# E-cadherin-mediated neighborhood surveillance dictates pre-malignant outcomes

**DOI:** 10.64898/2026.09.02.748765

**Authors:** Elizabeth A.N. Thompson, Clara Schimmer, Tatiana Omelchenko, Michele Paganini, Jesse S.S. Novak, Sienna T. Muller, Alain R. Bonny, Marina Schernthanner, Sairaj M. Sajjath, Charlotte J. Bell, Alp Gorgulu, S. Martina Parigi, Katherine S. Stewart, Priyam Banerjee, Elaine Fuchs, Stephanie J. Ellis

## Abstract

Stratified epithelia accumulate oncogenic mutations throughout life, yet overgrowths are rare. How epithelia detect and eliminate aberrant clones remains poorly understood. Using a mouse model of oncogenic clonal mosaicism in the skin, we find that pre-malignant epidermal cells redistribute E-cadherin to interfaces shared with wild-type neighbors, generating local tension heterogeneity that triggers elimination by cell competition. We show that gain or loss of E-cadherin can each drive competitive elimination, and although mechanical routes differ, both establish tension heterogeneity between neighbors, rather than any absolute adhesion state, as the critical determinant of epidermal fitness. This mechanism carries the seeds of its own failure: these tension differentials precipitate clonal sorting, depleting the wild-type contacts that surveillance requires. Pre-malignant cells then become supercompetitors, eliminating wild-type neighbors and expanding hyperplastically. Mechanical heterogeneity therefore endows tissues with an active, yet inherently fragile error-correction system whose collapse initiates a switch in competitive status, increasing susceptibility to tumorigenesis.

## Main

Over lifetimes, epithelial tissues become increasingly mosaic due to the inevitable accumulation of genetic mutations(*1*). Particularly in highly mutagen-exposed barrier tissues such as skin and esophagus, as many as 25% of cells from disease-free tissue have been reported to carry driver mutations for squamous cell carcinoma (SCC), one of the most common and potentially life-threatening cancer world-wide (*2, 3*). Yet, even in the face of these “pre-malignant” residents, an epithelium often maintains homeostasis.

How healthy and pre-malignant cells can harmoniously co-exist within tissues is poorly understood, as is the molecular trigger that tips the balance to drive cellular overgrowth and tumor initiation. A conceptual framework to understand this process comes from landmark cell competition studies in *Drosophila*, in which phenotypically weak “loser” cells are recognized and eliminated by more fit “winner” neighbors(*4–6*). In many well-studied models, including Myc-driven and other genetic paradigms, competitive status is tied to differences in proliferative capacity. Yet, competition can also arise between populations that are proliferatively matched, pointing to fitness signals that are not purely growth-based, and instead are rooted in aberrant structural features. Tellingly, in *Drosophila,* small clones of the cell polarity gene *scribble* behave as losers, yet can also drive overgrowth (*7–9*), suggesting the possibility that winner/loser status may not be set by genotype, but by tissue context. This notion might have relevance for pre-malignancy in mammalian tissues, where mosaic expression of oncogenes or tumor suppressor mutations can lead to either their elimination or homeostatic maintenance within the epithelial sheet (*10–16*), even though such mutations have the capacity to contribute to tissue overgrowth(*17–21*). Despite profound implications for cancer, the mechanisms by which the same mutation can confer loser or winner status depending on neighborhood composition, and how competitive status can change within a tissue, remain largely unresolved.

Emerging evidence points to cellular mechanics as a key determinant of competitive outcome(*22, 23*). Super-competitive cells can exploit competitive interactions to drive tumorigenic expansion by maximizing heterotypic contact with wild-type neighbors(*24, 25*), pointing to the importance of physical cellular interactions in determining fate. More broadly, differences in interfacial tension at clonal boundaries between wild-type and aberrant cells have been shown to drive actomyosin reorganization and loser cell elimination across several biological contexts(*26–29*). Cell-cell adhesion proteins have emerged as central candidate mediators of such mechanical interactions, but through mechanisms still largely unclear(*29–32*). Whether intercellular adhesion and/or tension differences between neighbors constitute the active mechanical fitness signal that drives context-dependent competitive outcome and whether and how quantitative differences in intercellular adhesion might be involved in pre-malignant growth control are questions that remain unaddressed.

Mammalian skin epidermis offers an ideal system to address these questions, and at the same time explore from the lens of oncogenic cell competition how a stratified squamous epithelium maintains tissue homeostasis. In contrast to fly epithelia, mammalian epidermis contains a monolayer of stem cells (EpSCs) that continuously self-renew and produce differentiated progeny(*33*) to form the stratified layers of the skin’s barrier. Moreover, in adult human skin, EpSCs can acquire oncogenic and tumor-suppressor mutations long before they disrupt normal homeostasis(*3*).

To tackle the early steps in premalignancy, and assess whether adhesion and mechanical heterogeneity might be at play in determining whether a potentially oncogenic cell will be eliminated or tolerated within the epidermis, we used a model in which we could induce SOX2 expression mosaically in developing mouse epidermis. SOX2 is a transcription factor not expressed by healthy EpSCs, but found in malignant, invasive SCCs of both mouse and human (*34*). Importantly, in mouse epidermis, ectopic SOX2 is both necessary and sufficient to drive hyperplasia, but on its own does not cause malignant invasion(*35, 36*)(Fig. 1A), making it an attractive model to interrogate how pre-malignant cells are normally restrained from launching oncogenic programs(*37, 38*).

**Figure 1.**
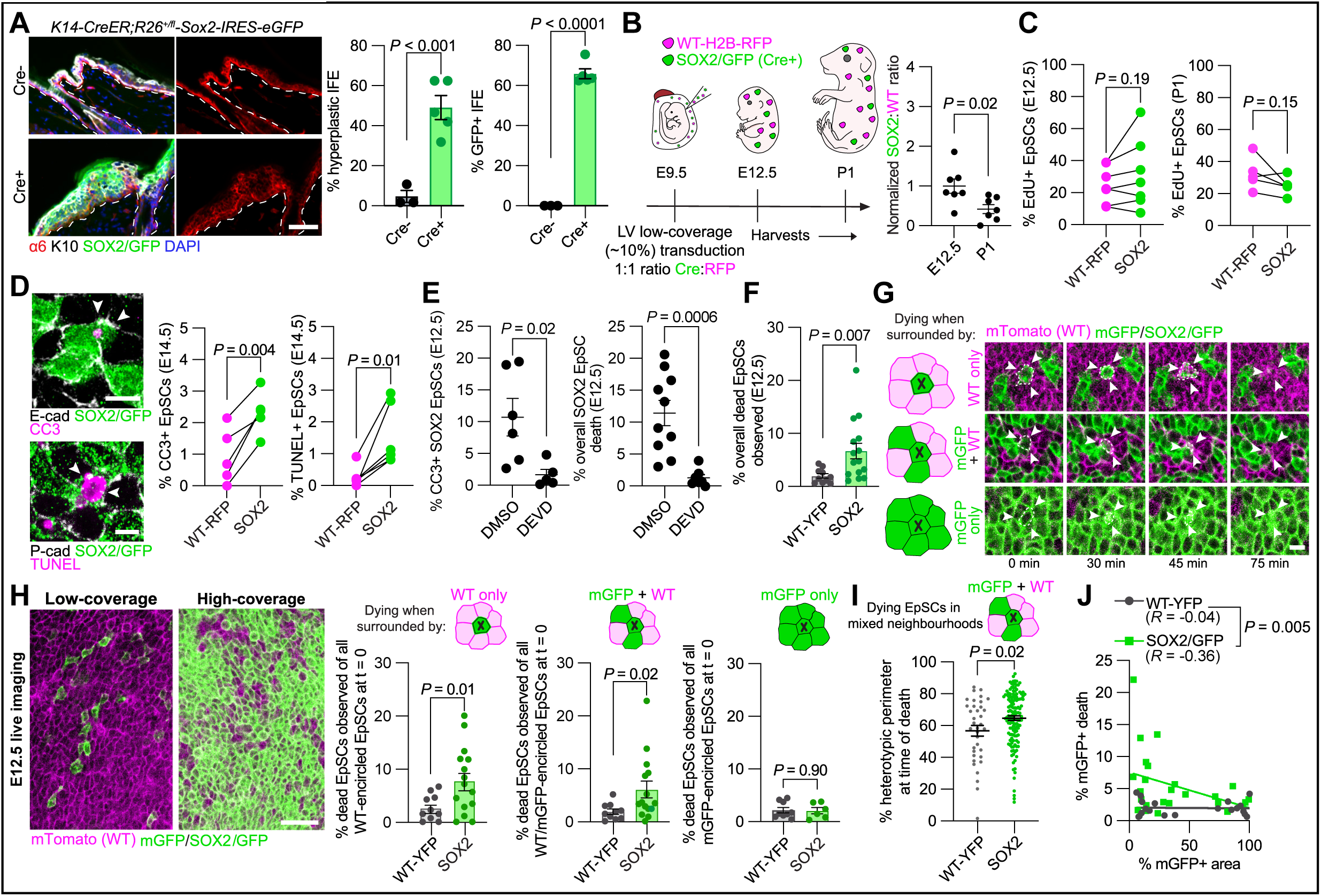
Cell competition drives elimination of sparse, pre-malignant SOX2 EpSCs. **A) Left**, representative sagittal immunofluorescence images taken from the backskins of *K14CreER*-*R26^+/fl^-Sox2-IRES-eGFP* animals (Cre- **top/**Cre+ **bottom**) treated with tamoxifen at second telogen (P49) and harvested 3 weeks post-tamoxifen. **Middle,** quantification of percentage of IFE that was hyperplastic, as defined as multi-layered α6 staining. **Right,** quantification of percentage of IFE staining positive for GFP. *n ≥* 3 mice per genotype; two-tailed unpaired *t-*test with Welch’s correction. **B)** Comparative growth assay (CGA) strategy reveals representation of SOX2 to WT EpSCs during embryonic development. Ratio of SOX2 (eGFP) to WT (RFP) EpSCs decreases over time. *n* ≥ 2 embryos per litter; 2 litters per time point; two-tailed unpaired *t*-test. **C)** EdU incorporation assays reveal that proliferation is comparable between SOX2 and WT EpSCs at E12.5 **(left)** and P1 **(right)**. *n* ≥ 2 embryos per litter; 2 litters per time point; two-tailed paired *t*-test. **D) Left,** whole-mount maximum projection immunofluorescence image of the mosaic E14.5 embryonic basal layer labelled for cleaved caspase-3 (**top**; CC3) and TUNEL (**bottom**) signals. **Right,** quantifications of the data, revealing signs of elevated death in SOX2 (white arrowheads) but not WT EpSCs. *n* ≥ 2 embryos per litter; 2 litters per time point; two-tailed paired *t*-test. **E) Left,** quantification of CC3+ SOX2 EpSCs in mosaic embryos treated with DMSO and caspase 3 inhibitor DEVD. *n* ≥ 2 embryos per litter; 2 litters per condition. **Right,** whole-mount live imaging analysis of dying SOX2 EpSCs in mosaic embryos treated with DMSO and DEVD. *n* ≥ 4 low-coverage FOVs per embryo, 2 embryos per conditio; two-tailed unpaired *t*-test with Welch’s correction. **F)** Quantification of total EpSC death events in low-coverage (competitive) mosaics between WT and SOX2 embryos. WT EpSCs were marked by either mTomato, or by YFP, from separate control experiments on *R26-lox-stop-lox-YFP* embryos transduced with LV-Cre (shown are WT data from the *R26-YFP* comparison, which was superior in that it controlled for the expression of Cre). Due to anatomical differences in LV transduction, individual 20X FOVs were used in lieu of whole embryos due to the vastly different neighborhood conformations yielded across the embryo (see **fig. S1E).** *n* = 9 FOVs across 3 embryos (WT-YFP); *n =* 15 FOVs across 5 embryos (SOX2); two-tailed unpaired *t*-test with Welch’s correction. **G)** Whole-mount live imaging analysis of representative dying E12.5 SOX2 EpSCs (dashed lines; arrowheads; black “X” on schematic) binned into one of three categories depending on neighbor contacts at time of death: **top row**, surrounded by WT (RFP) EpSCs only; **middle row**, in contact with both WT (RFP) and SOX2 (membrane mGFP and cytoplasmic eGFP) EpSCs; and **bottom row**, surrounded by SOX2 EpSCs only. **H)** Representative FOVs of low-coverage **(left)** and high-coverage **(right)** SOX2 mosaics and accompanying quantifications comparing WT-YFP and SOX2 EpSC death; *n =* 8 20X fields of view (FOVs) across 2 embryos (WT-YFP); 5 FOVs across 3 embryos (SOX2). **I)** Perimeter analysis of heterotypic contact performed between dying WT-YFP and SOX2 EpSCs found in mixed neighborhoods (contacting mGFP+ and mT+ neighbors). *n =* 36 WT-YFP and 138 SOX2 EpSCs; Mann-Whitney U test. **J)** Correlation analysis comparing mGFP+ area of the tissue and the corresponding amount of mGFP+ EpSC death observed. Includes intermediate-titer/coverage FOVs. *n* = 19 FOVs across 5 embryos (YFP); 24 FOVs across 9 embryos (SOX2). Statistics performed on nonlinear regressions. All data shown as mean ± s.e.m. Scale bars = 50 µm **(A, H)** and 10 µm **(D, G)**.

Using this model, we now identify E-cadherin as a master post-translational regulator of cell competition in the epidermis. SOX2 does not alter absolute expression levels of E-cadherin in EpSCs; rather, upon contact with wild-type neighbors, E-cadherin redistributes to heterotypic borders, generating the intercellular tension differences that determine competitive outcome. Manipulating E-cadherin levels in either direction is sufficient to trigger elimination through mechanically distinct routes, revealing a bidirectional logic of quality control. This surveillance is inherently fragile: as pre-malignant clones expand, the same tension differences that mark them for elimination drive their coalescence, eroding the heterotypic contacts on which surveillance depends. Beyond a critical clonal density, pre-malignant cells shed their loser identity and begin to eliminate wild-type neighbors. We thus unearth a transition from tumor suppression to tumor initiation driven not by new mutations, but by a shift in neighborhood composition, explaining why, in an intact mammalian tissue, a single oncogenic genotype can behave as either winner or loser through a mechanical, proliferation-independent competition.

### Cell competition drives elimination of sparse, pre-malignant SOX2 EpSCs

To interrogate cell-cell interactions between healthy and pre-malignant SOX2 EpSCs, we used *in utero* lentivirus (LV)-mediated delivery to specifically and stably transduce the early progenitors of embryonic day 9.5 (E9.5) ectoderm(*39*). Delivering LV-Cre and LV-H2B-RFP at low multiplicity of infection (MOI) into *R26-lox-stop-lox-Sox2-IRES-eGFP* embryos generated mosaic clones of SOX2 (eGFP+) and WT EpSCs (H2B-RFP+). Three days post-infection, the ratio of transduced GFP: RFP cells in E12.5 epidermis was ∼1.0. By postnatal day 1 (P1), this ratio had declined to 0.5:1.0 (Fig. 1B, fig. S1A). Analogous transductions in control *R26-lox-stop-lox-YFP* animals preserved the 1:1 YFP:RFP ratio over developmental time.

It seemed unlikely that as a root cause of hyperplasia, SOX2 would cause reduced proliferation. Indeed, EdU incorporation assays confirmed that the decreasing GFP:RFP ratio was not due to a proliferation difference (Fig. 1C, fig. S1B). Rather, more GFP+SOX2 EpSCs were positive for markers of apoptosis (cleaved caspase 3; CC3) and dying cell corpses (TUNEL) than their RFP+ EpSC counterparts, which served as internal WT controls within each embryo (Fig. 1D). Moreover, culturing embryos in the presence of caspase 3 inhibitor Z-DEVD-FMK (hereafter called DEVD) was also sufficient to prevent SOX2 EpSC death (Fig. 1E, fig. S1C).

Our findings thus far were consistent with the notion that SOX2 EpSCs may be actively eliminated when confronted with WT neighbors. To test whether the elimination of SOX2 EpSCs was dependent upon direct contact with WT EpSCs, we turned to live imaging of the epidermis of *R26^mTmG^;R26-lox-stop-lox-Sox2-IRES-eGFP* E12.5 embryos transduced at E9.5 with LV-Cre (fig. S1D). By delineating the borders of WT (mTomato) and SOX2 (mGFP, eGFP cytoplasm) EpSCs, we visualized each dying EpSC in real-time and in the context of its surrounding cellular neighborhood(*40*).

Using morphological features to characterize apoptosis, with LV-Cre transduced *R26^mTmG^;R26-lox-stop-lox-YFP* embryos as controls, we found that mGFP+ SOX2 EpSCs died more frequently than YFP+ WT EpSCs in mosaic embryos (Fig. 1F, fig. S1E). By varying viral titer to generate regions of low and high SOX2 coverage, we examined apoptotic events across a full spectrum of neighborhoods. Binning dying mGFP+ EpSCs by their neighborhood contacts, we found that SOX2 EpSCs died primarily when contacting WT (mTomato+) neighbors (Fig. 1, G and H, fig. S1F, Supplementary Videos S1-S3). Within mixed neighborhoods, and in contrast to WT-YFP controls, dying SOX2 EpSCs made more extensive contacts with WT than with SOX2 EpSCs, and SOX2 EpSCs showed a striking inverse correlation between their tissue representation and their degree of death (Fig. 1, I and J, fig. S1G). Together, these data establish that SOX2 EpSC cells are eliminated by cell competition, behaving as losers, but only when outnumbered by WT neighbors. The data further suggested that direct heterotypic cell-cell contact, rather than SOX2-induced cell autonomous differences in cell proliferation rate, is the critical determinant of loser fate. We substantiate this hypothesis further below.

We next asked how contact-dependent apoptosis relates to SOX2 EpSC behavior across the full window of epidermal expansion. Reminiscent of our prior studies on *Mycn*-mediated cell competion(*40*), SOX2 clones at later developmental stages exhibited an increased propensity to differentiate, contributing additionally to their loser phenotype and loss from the basal layer (Fig. S1H). To further assess the relative contribution of each cellular behavior across development, we developed a statistical model incorporating measured rates of proliferation, apoptosis, and differentiation in their respective windows of activity (fig. S2, A and B, see Methods).

The model recapitulated the observed SOX2:WT trajectory from E12.5 to P1 (fig. S2, B to F). Furthermore, phase decomposition revealed that early apoptosis (E12.5–E15.5) and late differentiation (E15.5–P1) operate in temporally distinct windows, together accounting for the full reduction in SOX2 EpSC representation observed at birth. Thus, we were able to examine single-cell elimination in the absence of the stochastic homeostatic flux of differentiation-driven basal-layer loss that emerged later by focusing on early development, where apoptosis was the sole driver of SOX2 loss and sparse SOX2 EpSCs made extensive contact with WT neighbors.

### SOX2 EpSC elimination correlates with increased intercellular tension at heterotypic contacts

To investigate how heterotypic contact marks SOX2 EpSCs for elimination and more readily pinpoint cellular properties underlying their loser phenotype, we analyzed SOX2 EpSCs entirely surrounded by WT neighbors — so-called ‘SOX2 islands’. SOX2 islands consistently exhibited increased cell area and junction length compared to both WT neighbors and to other SOX2 EpSCs in the basal epidermis at E14.5 (Fig. 2A) suggesting a non-cell-autonomous effect of the local neighborhood on cell size and junction length. This led us to posit that SOX2 islands may be under tension, effectively being stretched by their WT neighbors.

**Figure 2.**
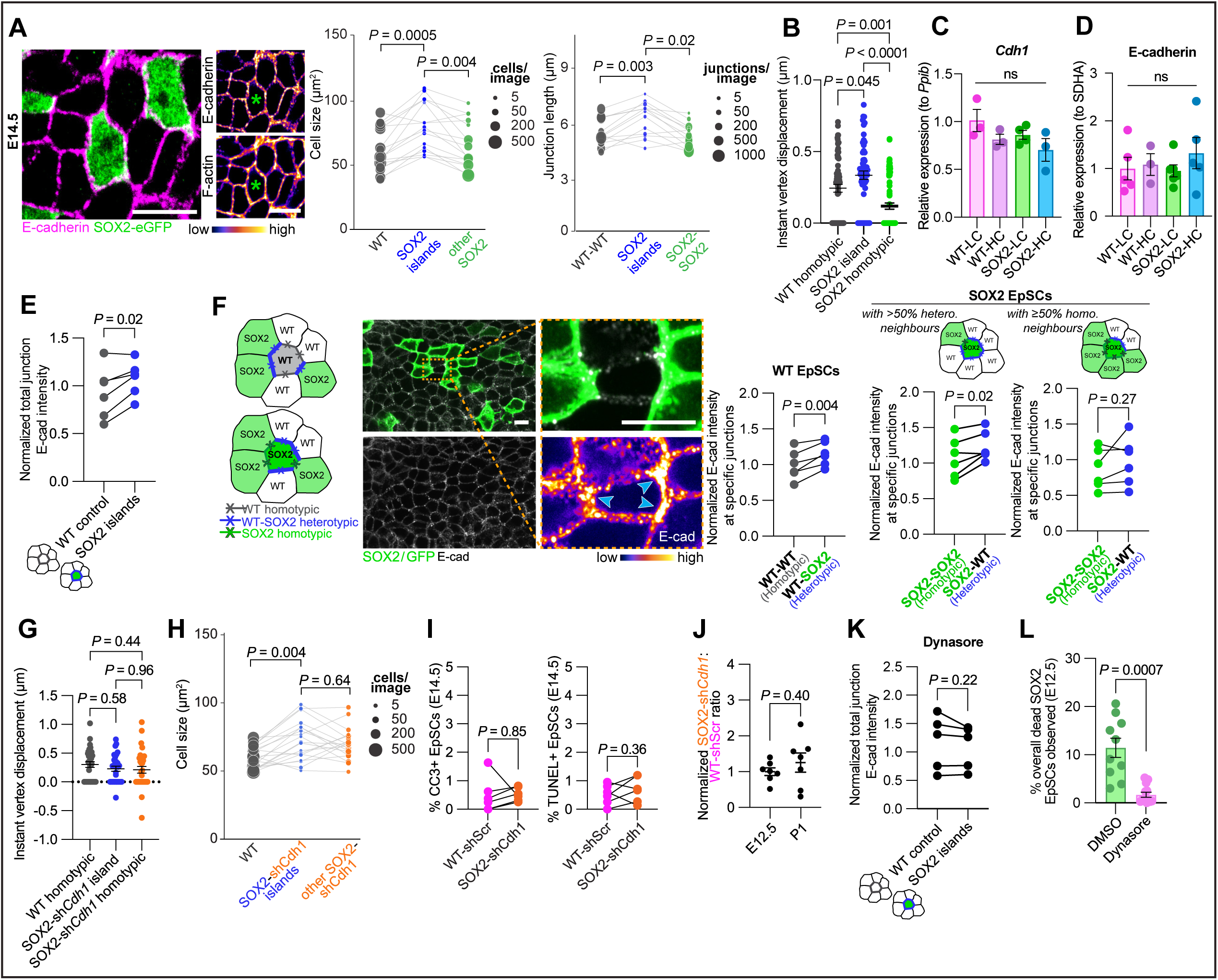
Mechanical and molecular analysis of the heterotypic SOX2-WT interface identifies post-translational redistribution of E-cadherin as a driver of mechanical competition. **A) Left,** whole-mount maximum projection immunofluorescence image of E14.5 SOX2 EpSC mosaics comprised of larger SOX2 “islands” stained with E-cadherin **(top)** and F-actin **(below)**. Green asterisks denote SOX2 island. **Middle,** quantification of cell size between WT controls, SOX2 islands, and all non-island SOX2 EpSCs. **Right,** quantification of individual lengths between WT-WT homotypic, WT-SOX2 heterotypic island, and SOX2-SOX2 homotypic junctions. For cell size and junction data, each datapoint represents measurements extracted from the average value of all cells in each category within a single FOV/image; lines connect matched values from within the same image. *n* = 16 images across 8 embryos from 4 litters. For cell size, comparisons between islands were made via sign tests; images that had <2 islands were excluded from statistical tests making the effective *n* = 14. Non-island categories, and all comparisons within the junction length dataset, were compared via a paired t-test with x3 Bonferroni correction. **B)** Quantification of instant vertex displacement following laser ablation in E14.5 embryos between WT-WT homotypic, WT-SOX2 island heterotypic, and SOX2-SOX2 homotypic junctions. Two vertices are plotted per junction. *n* = 36 WT-WT homotypic; 30 WT-SOX2 heterotypic islands; 42 SOX2-SOX2 homotypic junctions (across ≥ 15 embryos, ≥ 8 litters total); one-way ANOVA with Tukey’s multiple comparisons test. **C)** qPCR and **D)** Western blot analyses on sorted E14.5 EpSCs comparing *Cdh1* mRNA transcripts (normalized to *Ppib*) **(C)** and E-cadherin protein (normalized to SDHA) **(D)** between sorted LC WT, HC WT, LC SOX2, and HC SOX2 EpSC populations. *n* ≥ 3 litters per condition; ≥ 2 embryos pooled per litter; one-way ANOVA with Tukey’s multiple comparisons test. **E)** Quantification of E-cad intensity across all junctions between control WT and SOX2 islands. Values are normalized to average WT control value across all embryos and litters. *n* ≥ 2 embryos per litter; 2 litters total; two-tailed paired *t*-test. **F) Far left,** schematic of homo/heterotypic border analyses. **Left,** representative whole-mount maximum projection immunofluorescence image of an E14.5 basal epidermis mosaically populated with SOX2 EpSCs (GFP+) and immunostained for E-cadherin. **Inset,** magnified WT EpSC with heterotypic (cyan arrowheads) and homotypic borders. **Right,** accompanying quantifications of normalized E-cadherin immunostaining between homotypic and heterotypic borders in WT (all neighborhood conformations) and SOX2 (with either >50% heterotypic or ≤ 50% homotypic neighbors) EpSCs. *n* = 3 embryos per litter; 2 litters total; two-tailed paired *t-*test. **G)** Quantification of instant vertex displacement following laser ablation in E14.5 embryos between WT-WT homotypic, WT-SOX2^sh*Cdh1*^ island heterotypic, and SOX2^sh*Cdh1*^-SOX2^sh*Cdh1*^ homotypic junctions. Two vertices are plotted per junction. *n* = 15 WT-WT homotypic; 16 WT-SOX2^sh*Cdh1*^ heterotypic islands; 16 SOX2^sh*Cdh1*^-SOX2^sh*Cdh1*^ homotypic junctions (across ≥ 7 embryos, ≥ 4 litters total); one-way ANOVA with Tukey’s multiple comparisons test. **H)** Quantification of cell size between WT controls, SOX2^sh*Cdh1*^ islands, and all non-island SOX2^sh*Cdh1*^ EpSCs. Each datapoint represents measurements extracted from the average value of all cells in each category within a single field of view; lines connect matched values from within the same image. *n* = 19 images across 5 embryos from 2 litters. Statistical comparisons between categories were made via paired *t*-tests with x3 Bonferroni correction. **I)** Analogous CC3 and TUNEL quantifications to Fig. 1D, but instead of comparing SOX2^shScr^ and WT^shScr^ EpSCs within the same embryo, now comparing SOX2^sh*Cdh1*^ and WT-RFP^shScr^*. n ≥* 2 embryos per litter, 2 litters total; two-tailed paired *t-*test. **J)** Comparative growth assay measuring representation of SOX2^sh*Cdh1*^ EpSCs to WT-RFP^shScr^ controls over development. *n ≥* 2 embryos per litter; 2 litters per time point; two-tailed unpaired *t*-test. **K)** Quantification of E-cadherin intensity across all junctions between control WT and SOX2 islands in Dynasore-treated embryos at E14.5. Values are normalized to average control WT across all DMSO-treated embryos and litters. *n* ≥ 2 embryos per litter; 2 litters total; two-tailed paired *t*-test. **L)** Quantification of SOX2 EpSC death in live-imaged E12.5 embryos treated with either DMSO or Dynasore. *n =* 1-5 FOVs per embryo; 3 embryos per condition; two-tailed unpaired *t*-test with Welch’s correction. All data shown as mean ± s.e.m. Scale bars = 10 µm.

Since mechanical stretch has recently been shown to be a potent trigger of the homeostatic villus cell shedding that occurs in the intestinal epithelium(*41*), we wondered if a similar process might be driving elimination of SOX2 EpSCs from the basal epidermis. To test this possibility, we used mTmG membrane fluorescence to mark each kind of intercellular contact and laser ablation to assess the consequences of severing heterotypic versus homotypic interfaces (Supplementary Videos S4-S6).

Proportional to junctional tension(*42*), instantaneous vertex displacement following ablation was significantly higher at heterotypic junctions than at either homotypic junction class (Fig. 2B, fig. S3A). Notably, SOX2/SOX2 junctions exhibited the smallest vertex displacement, suggesting that they are under less tension than WT/WT junctions even at baseline. Intriguingly, when we examined heterotypic junctions across a broader range of SOX2 EpSCs with varying neighborhood compositions, we found that retraction displacement scaled with the proportion of cell perimeter shared with wild-type neighbors: the more wild-type contact, the greater the recoil (fig. S3B). These data pointed to a dose-dependent effect of competitive context on junctional tension.

Beyond instantaneous retraction, time-lapse imaging of junction relaxation over 50–60 seconds after ablation allowed us to fit the data to a viscoelastic model(*42*) and extract force-to-elasticity ratios (*D*) and relaxation half-times (tau) for each junction class. Force-to-elasticity ratios were significantly higher at heterotypic WT-SOX2 junctions than at either homotypic class, suggesting greater forces acting across the competitive interface (fig. S3C). The force-to-elasticity ratios of homotypic junctions did not differ from one another, supporting the conclusion that elevated tension was a specific product of heterotypic neighbor interactions rather than a cell-autonomous property.

Relaxation half-times, which reflect tissue viscosity, gave additional insight. SOX2-SOX2 junctions exhibited longer tau than WT-WT junctions, implying that SOX2 EpSCs were intrinsically more viscous — a cell-autonomous mechanical property that also aligned with the smaller instantaneous retraction displacements (Fig. 2B, fig. S3C). By contrast, tau at heterotypic junctions was indistinguishable from either homotypic class, indicating that viscosity, unlike tension, was not modulated by neighbor identity. Thus, distinct mechanical properties — in particular elevated intercellular tension — emerged specifically at the competitive WT-SOX2 interface.

### E-Cadherin mediates differential SOX2 EpSC mechanics and loser status

We next sought to understand the molecular basis of the atypical mechanical interfaces at heterotypic borders. We began by focusing on E-cadherin, as it is known to form the transmembrane core of the intercellular adherens junctions that associate with the actin cytoskeleton and function in mechanotransduction within the EpSC monolayer(*43–45*). However, at the RNA and protein levels, we found no differences in E-cadherin levels in EpSCs isolated from E14.5 SOX2 versus WT mouse skin (Fig. 2, C and D, fig. S3, D and E). Moreover, cell neighbor status did not influence these levels, which were the same whether the density of SOX2 EpSCs in mosaic skin was high (non-competitive) or low (competitive) relative to WT EpSCs (shown). Similarly, when we quantified E-cadherin immunofluorescence in non-competitive SOX2 and WT regions of E14.5 SOX2 mosaic epidermis, we observed no differences in total levels of junctional E-cadherin across genotypes (fig. S3E). Thus, in this context, the SOX2 oncogene did not appear to affect E-cadherin expression nor the competitive status of SOX2.

In striking contrast, when we examined boundaries of SOX2 and WT clones, we found that when surrounded by WT EpSCs, SOX2 islands exhibited significantly elevated total junctional E-cadherin compared to WT control EpSCs (Fig. 2E). Moreover, enrichment was restricted to heterotypic junctions: E-cadherin intensity at SOX2-WT junctions was consistently higher than at SOX2-SOX2 homotypic junctions within the same cell (Fig. 2F). This effect was also contact-dependent, since SOX2 EpSCs surrounded predominantly by WT neighbors showed clear increases in E-cadherin intensity at heterotypic junctions, whereas SOX2 EpSCs with predominantly homotypic neighbors did not. Notably, the enrichment was bilateral: E-cadherin was also elevated at heterotypic junctions relative to homotypic junctions within WT EpSCs (shown), revealing an intriguing asymmetry in E-cadherin distribution between competitive and non-competitive interfaces.

Two critical control experiments confirmed that this enrichment depended specifically on competitive WT-SOX2 interactions. First, non-competitive SOX2 EpSCs (i.e., SOX2 EpSCs in tissue dominated by other SOX2 EpSCs) and WT EpSCs exhibited equivalent total junctional E-cadherin intensities, ruling out a baseline difference in E-cadherin expression between the two cell populations (fig. S3, D and E). Second, in WT-YFP mosaic embryos, no significant differences in E-cadherin intensity were detected at heterotypic versus homotypic borders (fig. S3, F and G), excluding the transgenic mosaic configuration itself as a trigger for redistribution. By contrast, this redistribution was accompanied by parallel reorganization of cortical F-actin at heterotypic junctions, consistent with the elevated tension we observed at heterotypic contacts (fig. S3H).

Our results thus far suggested that E-cadherin-mediated mechanical competition underlies the differential SOX2 EpSC behaviors and elimination by WT neighbors. If so, depleting E-cadherin in SOX2 EpSCs should rescue these differences and SOX2 EpSC elimination in a competitive environment. To functionally test this prediction, we used RNA interference and confirmed a previously validated short *Cdh1* hairpin(*46*) to effectively reduce E-cadherin specifically in SOX2 EpSC clones (fig. S4, A and B).

We first used laser ablation to assess how *Cdh1* knockdown affects junctions in SOX2*^shCdh1^* mosaic epidermis at E14.5. In contrast to scramble controls, the instantaneous retraction displacements previously seen at SOX2:WT interfaces were indistinguishable from WT:WT interfaces upon selective depletion of E-cadherin in SOX2 cells (Fig. 2G). Parameter fitting reinforced this finding: heterotypic junctions no longer displayed different force-to-elasticity ratios or relaxation half-times (fig. S4C). These data supported the view that E-cadherin is required to generate the distinct mechanical signature of the competitive interface.

*Cdh1* depletion did produce cell-autonomous changes in SOX2-SOX2 junctions, further reducing force-to-elasticity ratios and indicating that E-cadherin loss has additional cell-intrinsic effects on the SOX2 mechanical phenotype. Critically, however, these intrinsic changes did not propagate to the heterotypic interface, where mechanical heterogeneity was abolished — confirming that the surveillance signal resided specifically at the competitive boundary rather than within either cell population.

Cell shape analysis revealed that although SOX2*^shCdh1^* EpSCs remained larger than WT cells, the additional, island-specific increase in cell area and junction length was lost (Fig. 2H, fig. S4D), further indicating that the neighborhood-dependent component of the phenotype was rescued by downregulation of E-cadherin in SOX2 EpSCs. Confirming rescue of loser status, SOX2*^shCdh1^* EpSCs and their internal WT-RFP controls showed comparable CC3 and TUNEL labelling (Fig. 2I), and live imaging confirmed that SOX2*^shCdh1^* EpSCs no longer matched the elevated apoptosis of SOX2^shScr^ EpSCs. Although SOX2^sh*Cdh1*^ mosaics showed a modest increase in total death events relative to WT-YFP controls, SOX2^sh*Cdh1*^ EpSCs in WT-only or mixed neighborhoods died at frequencies indistinguishable from WT-YFP controls (fig. S4E). CC3 labelling was also unchanged in high-coverage SOX2^sh*Cdh1*^ regions relative to SOX2^shScr^ controls, indicating that baseline apoptotic sensitivity was unaltered by *Cdh1* knockdown (fig. S4F). Finally, comparative growth assays showed that SOX2^sh*Cdh1*^ EpSCs were retained and grew comparably to controls over development (Fig. 2J).

Although depleting E-cadherin in SOX2 EpSCs erased their loser status, it did not convert them into winners (fig. S4G). These data argue against a simple adhesion-asymmetry model of competition based purely upon differential E-cadherin levels at heterotypic borders. Consistent with this interpretation, P-cadherin, which can compensate for E-cadherin in maintaining epidermal adhesion(*43*), was upregulated in SOX2^sh*Cdh1*^ EpSCs, and yet rescue still occurred, further dissociating adhesion status from competitive outcome (fig. S4B). Rather, our collective data provided compelling evidence that it is tension heterogeneity at heterotypic interfaces, specifically mitigated upon *Cdh1* depletion, that constitutes the critical fitness signal.

Given E-cadherin’s importance, what are the underlying mechanisms that regulate its enhanced localization to winner:loser borders? Since E-cadherin levels were not affected by genotype nor the non-competitive versus competitive environment of SOX2 EpSCs, our results pointed instead to the view that E-cadherin may be dynamically redistributed upon heterotypic contact. One potential mode of redistribution is through endocytic recycling of E-cadherin at the cell surface.

To test this possibility, we cultured sparse SOX2 mosaic E14.5 embryos in the presence of Dynasore, a Dynamin GTPase inhibitor widely used to block endocytic protein dynamics(*47*). Dynasore treatment resulted in reduced E-cadherin at all cell-cell junctions, consistent with a general requirement for endocytosis in maintaining junctional E-cadherin (fig. S4H). Dynasore treatment also equalized E-cadherin levels between SOX2 islands and WT controls (Fig. 3K) and abolished the differential retraction dynamics at heterotypic versus homotypic interfaces (fig. S4I). Most strikingly, apoptosis in SOX2 EpSCs was markedly reduced upon culturing E12.5 mosaic embryos in the presence of Dynasore (Fig. 2L). Together, these data are consistent with the notion that endocytosis-mediated E-cadherin redistribution is a key component of the mechanism underlying competitive elimination of SOX2 EpSCs.

**Figure 3.**
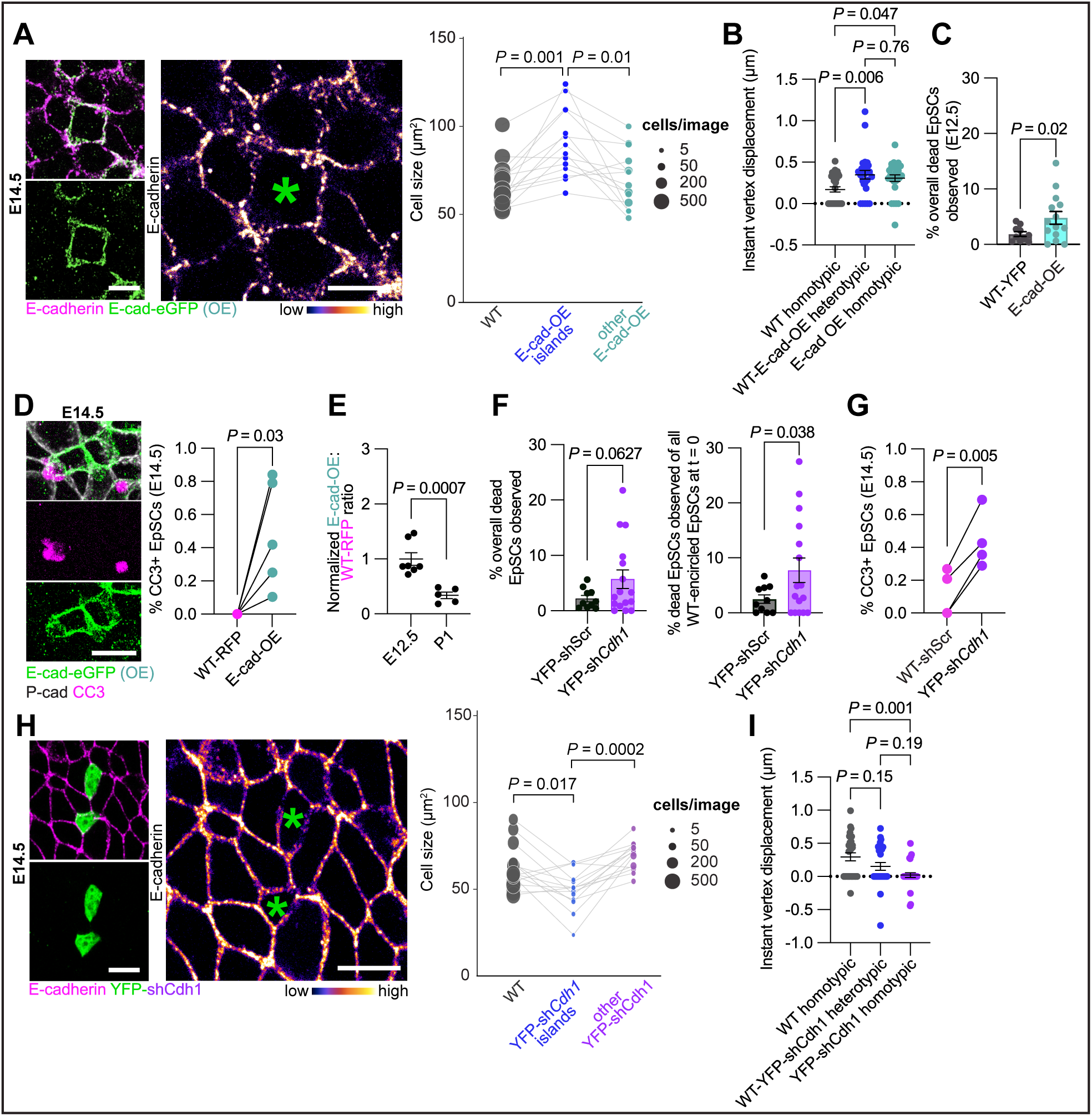
E-cadherin levels bidirectionally control competitive cell fate through mechanically distinct routes of elimination. **A) Left,** representative whole-mount maximum projection immunofluorescence image of E14.5 epidermis mosaically over-expressing E-cad-eGFP. Green asterisk denotes E-cad^OE^ island. **Right,** quantification of cell size between WT controls, E-cad^OE^ islands, and all non-island E-cad^OE^ EpSCs. Each datapoint represents measurements extracted from the average value of all cells in each category within a single field of view; lines connect matched values from within the same image. *n* = 14 images across 7 embryos from 2 litters. Comparisons between islands were made via sign tests. Non-island categories were compared via a paired *t*-test with Bonferroni correction. **B)** Quantification of instant vertex displacement following laser ablation in E14.5 embryos between WT-WT homotypic, WT-E-cad^OE^ heterotypic, and E-cad^OE^-E-cad^OE^ homotypic junctions. Two vertices are plotted per junction. *n* = 17 WT-WT homotypic; 13 WT-E-cad^OE^ heterotypic; 16 E-cad^OE^-E-cad^OE^ homotypic junctions (across ≥ 5 embryos, ≥ 4 litters total); one-way ANOVA with Tukey’s multiple comparisons test. **C)** Quantifications of death events from E12.5 live imaging experiments between WT-YFP controls (re-plotted from Fig. 1F) and E-cad^OE^ EpSCs, *n* ≥ 3 FOVs per embryo, 3 embryos per condition; two-tailed unpaired *t*-test with Welch’s correction. **D) Left,** representative whole-mount max projection immunofluorescence image of CC3+ E-cad^OE^ EpSCs in E14.5 mosaic epidermis and **right,** accompanying quantification comparing CC3 expression between E-cad^OE^ and WT controls. *n* ≥ 2 embryos per litter; 2 litters total; two-tailed paired *t*-test. **E)** Comparative growth assay measuring representation of E-cad^OE^ EpSCs to WT-RFP controls over the course of development. *n ≥* 2 embryos per litter; 2 litters per time point; two-tailed unpaired *t*-test. **F)** Quantifications of death events from E12.5 live imaging experiments between WT-YFP^shScr^ (re-plotted from Fig. 1F, H) and YFP^sh*Cdh1*^ EpSCs. Shown are percentages of EpSC deaths for: **left**, total; **right**, those encircled by WT EpSCs only. *n =* 2-5 FOVs per embryo; 3-4 embryos per condition; two-tailed unpaired *t*-test with Welch’s correction. **G)** Quantification comparing CC3 expression between WT-YFP^shScr^ and YFP^sh*Cdh1*^ EpsCs at E14.5. *n* ≥ 2 embryos per litter; 2 litters total; two-tailed paired *t*-test. **H) Left,** representative whole-mount maximum projection immunofluorescence image of E14.5 epidermis mosaically populated by YFP^sh*Cdh1*^ EpSCs. Green asterisks denote YFP^sh*Cdh1*^ islands. **Right,** quantification of cell size between WT controls, YFP^sh*Cdh1*^ islands, and all non-island YFP^sh*Cdh1*^ EpSCs. Each datapoint represents measurements extracted from the average value of all cells in each category within a single field of view; lines connect matched values from within the same image. *n* = 14 images across 3 embryos from 2 litters; paired t-test with 3x Bonferroni correction for multiple comparisons. **I)** Quantification of instant vertex displacement following laser ablation in E14.5 embryos between WT-WT homotypic, WT-YFP^sh*Cdh1*^ heterotypic, and YFP^sh*Cdh1*^-YFP^sh*Cdh1*^ homotypic junctions. Two vertices are plotted per junction. *n* = 14 of each junction (across ≥ 3 embryos, ≥ 2 litters total); one-way ANOVA with Tukey’s multiple comparisons test. All data shown as mean ± s.e.m. Scale bars = 10 µm.

To test whether elevating E-cadherin at heterotypic junctions is sufficient to drive loser fate, we generated epidermis containing a small proportion of EpSCs that overexpressed an E-cadherin-GFP fusion protein (hereafter referred to as E-cad^OE^), previously shown to behave similarly to its wild-type counterpart(*48*) (fig. S5, A and B). However, when E-cad^OE^ EpSCs were surrounded by wild-type neighbors, they exhibited larger cell area and increased instantaneous vertex displacement, the latter indicating elevated junctional tension, and the larger area consistent with cell stretching (Fig. 3, A and B, fig. S5, C and D). Parameter fitting corroborated the heterotypic-specific tension increase: force-to-elasticity ratios at heterotypic junctions were higher than homotypic wild-type junctions. In this regard, E-cad^OE^ EpSCs behaved like SOX2 EpSCs.

Functionally, live imaging of mosaic E12.5 embryos revealed that E-cad^OE^ EpSCs also died more often than control WT-YFP EpSCs in low-coverage mosaics (Fig. 3C). This result was confirmed at E14.5 by CC3 staining, where, within the same embryo, E-cad^OE^ EpSCs more frequently were positively labelled for CC3 than their RFP-labelled WT counterparts (Fig. 3D). Moreover, compared to WT EpSCs, E-cad^OE^ EpSCs exhibited an overall growth disadvantage at birth (Fig. 3E) and yet proliferation was not affected (fig. S5E). Overall, these data demonstrated that sparse E-cad^OE^ EpSCs displayed a loser phenotype during epidermal development and recapitulated the key features of SOX2 EpSC competitive elimination.

Our findings further indicated that wild-type EpSCs are sensitive to E-cadherin excess in neighboring cells. Interestingly, however, the epithelium was equally sensitive to E-cadherin deficit; thus, in otherwise wild-type tissue, clones of E-cadherin deficient EpSCs behaved as losers and underwent neighborhood-dependent apoptosis, as confirmed through both live imaging and cleaved caspase-3 immunofluorescence (Fig. 3, F and G). Further suggestive of a loser phenotype, sparsely induced YFP*^shCdh1^*EpSCs exhibited a trend pointing to a comparative growth disadvantage compared to WT EpSCs in epidermal development (fig. S5F). Apoptosis appeared to be the major driver of these growth dynamics at E12.5, as EdU incorporation was comparable between control and *Cdh1* deficient cells (fig. S5G).

In contrast to SOX2 and E-cad^OE^ islands, YFP*^shCdh1^*islands displayed reduced cell size (Fig. 3H). Laser ablation experiments also revealed decreases in force-to-elasticity ratios at heterotypic junctions compared to WT-WT homotypic junctions (fig. S5H). These data indicated that E-cadherin loss drove elimination through a mechanically distinct route, accompanied by cell compression rather than stretching. Consistently, instantaneous vertex displacement showed a graded trend — highest at WT-WT junctions, reduced at heterotypic junctions, and lowest at YFP*^shCdh1^*-YFP*^shCdh1^* junctions (Fig. 3I) — with a greater fraction of YFP*^shCdh1^* homotypic measurements registering negative displacements. Importantly, these data bolstered the link with mechanical tension that we had discovered when otherwise wild-type E-cadherin levels were redistributed asymmetrically in a competitive setting. Collectively, our data provided compelling evidence that E-cadherin acts as a bidirectional reporter of mechanical fitness, triggering elimination of cells that deviate too far in either direction from a tolerable homeostatic range.

### Tissue-scale properties of SOX2 mosaics enable evasion of tumor suppressive cell competition

Theoretical and experimental work in *Drosophila* indicate that differences in interfacial tension at clonal boundaries can drive cell sorting and shape clonal evolution over time(*28, 49*). Given the striking tensile differences we observed at heterotypic WT-SOX2 interfaces, we asked whether analogous mechanics might drive sorting of SOX2 clones from their WT neighbors during epidermal development — and whether this creation of larger clones has consequences for the pre-malignant surveillance mechanism we have described.

To explore this question, we first mapped our laser ablation onto a phase diagram that predicts clonal shape purely on the basis of the relative differences in tension at clonal interfaces(*49*) in wild-type or SOX2-expressing clones. Consistent with our expectation, this approach predicted that SOX2 clones would be more clustered and have a more rounded morphology than wild-type clones, which would be more dispersed (fig. S6A). Furthermore, the model predicted that SOX2^sh*Cdh1*^ clones should be less rounded and have a near WT-clonal shape.

To directly test these predictions, we transduced E9.5 *R26-lox-stop-lox-Sox2-IRES-eGFP* epidermis with low-titer LV-Cre or LV-H2B-RFP to ensure examination of clonal offspring derived from single SOX2 or WT EpSCs at E14.5. As shown in Fig. 4A, while the morphology of WT (H2B-RFP+) EpSC clones was dispersed, SOX2 clones were clustered. However, when we changed SOX2 clonal morphology to a non-competitive setting by altering the LV-Cre titer, SOX2 clones now closely resembled the dispersed morphology of wild-type control clones.

**Figure 4.**
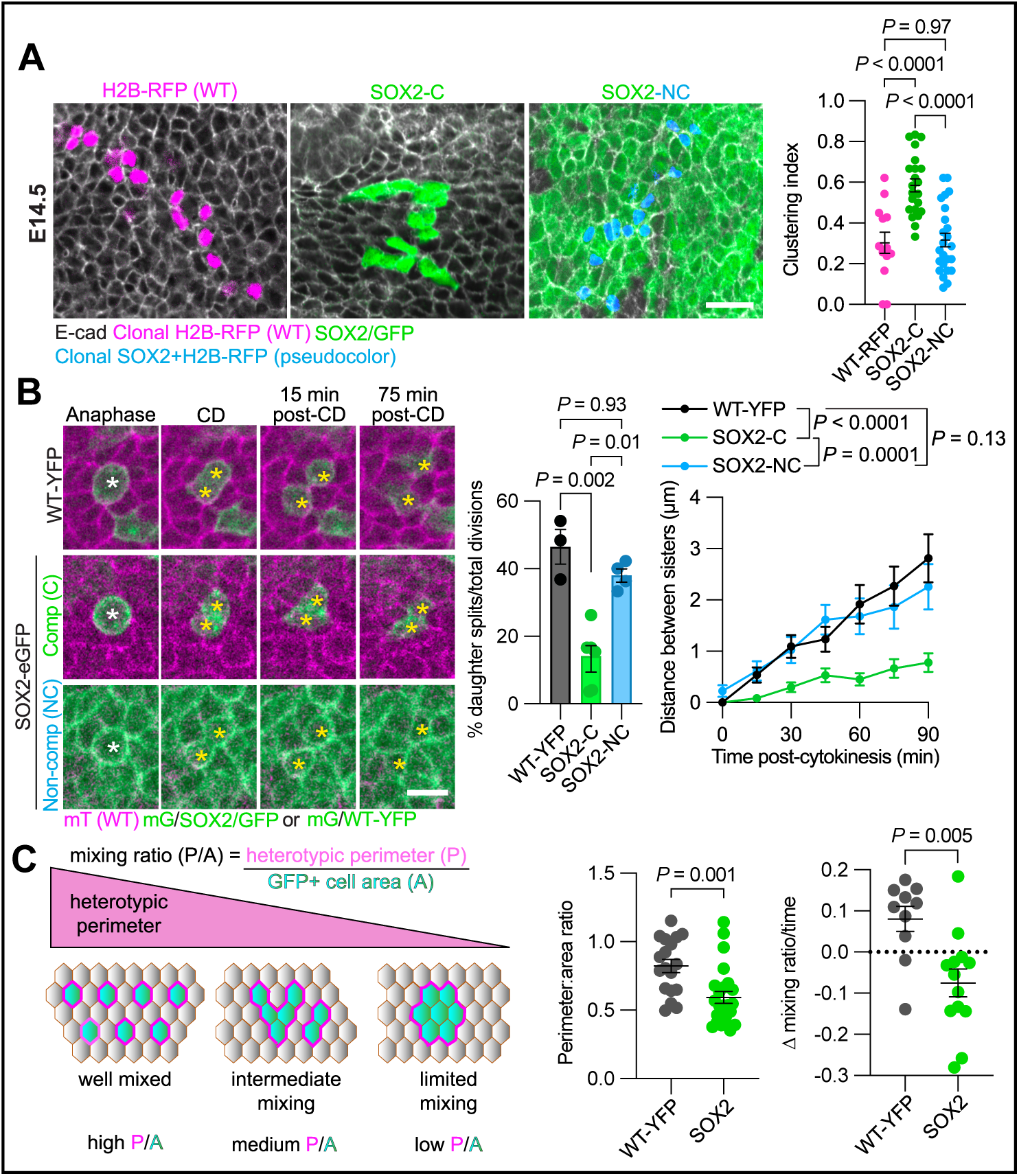
Tension-driven clonal sorting progressively reduces heterotypic contact in SOX2 mosaics. **A) Left,** representative whole-mount maximum projection immunofluorescence images of WT, SOX2 (competitive), and SOX2 (non-competitive) EpSC clones at E14.5. **Right,** quantifications of clustering behavior (see Methods and **fig. S6B**). For each condition, *n* ≥ 17 clones from 3-6 embryos across 2 litters; ordinary one-way ANOVA with Tukey’s multiple comparisons test. **B) Left,** E12.5 live imaging of representative dividing EpSCs, illustrating the differences in dispersion between SOX2 EpSC daughters post-cell division (post-CD) when they are in competitive **(middle row)** versus non-competitive **(bottom row)** neighborhoods. In non-competitive neighborhoods, SOX2 and WT EpSCs behave similarly (**bottom** and **top rows**, respectively). **Middle,** quantifications showing the percentages of EpSC clones that have dispersed within 45 min of CD. *n* = 3 embryos (WT-YFP); 6 embryos (competitive SOX2); 4 embryos (non-competitive SOX2); one-way ANOVA with Tukey’s multiple comparisons test. **Right**, quantifications at 15 min intervals post-division to plot the distance between the daughters. *n* = 73 WT EpSCs (3 embryos); 117 competitive SOX2 EpSCs (7 embryos); 66 non-competitive SOX2 EpSCs (5 embryos). Statistics performed on nonlinear regressions. **C)** Tissue-scale de-mixing of SOX2 EpSC clones quantified by perimeter-to-area mixing ratio. **Left,** schematic shows the P/A ratio: heterotypic contact perimeter (P) normalized to total labelled cell area (A). **Middle,** P/A in WT-YFP versus SOX2 mosaics at E12.5. **Right,** change in P/A over 12 hr of live imaging; negative values indicate progressive demixing. *n* ≥ 10 FOVs from 3 embryos (WT-YFP) and 15 FOVs from 5 embryos (SOX2); two-tailed unpaired *t*-test. All data shown as mean ± s.e.m. Scale bars = 20 µm **(A)**; 10 µm **(B)**.

Two quantitative measures — a clustering index based on labelled neighbor contacts, and a clonal dispersion assay (fig. S6, B and C) — confirmed that wild-type and non-competitive SOX2 clones grew dispersed, whereas sparse SOX2 clones surrounded by wild-type competitors grew more clustered. Clonality was verified using a *R26-lox-stop-lox-Brainbow2.1* model(*50*) (fig. S6D). Analogous analyses at E12.5 showed that SOX2 clones were already more cohesive than wild-type clones (fig. S7A), indicating that tension-driven sorting is an early and persistent feature of epidermal clonal dynamics. This clustering was E-cadherin-dependent, as SOX2^sh*Cdh1*^ clones were significantly more dispersed than SOX2^shScr^ clones (Fig. 4B, fig. S7B). Together, these data established that clonal clustering of SOX2 EpSCs depends on both neighborhood context and E-cadherin.

Next, to obtain direct dynamic evidence for tension-driven sorting, we turned to time-lapse imaging of dividing EpSCs in E12.5 embryos. In wild-type clones, daughter EpSCs separated substantially following mitosis (Fig. 4B, fig. S8A, Supplementary Video S7). In striking contrast, SOX2 daughter EpSCs remained in close proximity following division, consistent with preferential cohesion between SOX2-SOX2 interfaces relative to SOX2-WT interfaces (Supplementary Video S8). We quantified this behavior both by centroid tracking and cell border analysis, confirming that daughter cell separation was significantly reduced in SOX2 clones compared to either wild-type controls or SOX2 EpSCs dividing in a non-competitive environment (Fig. 4B, fig. S8A, Supplementary Video S9). We also found that this behavior was E-cadherin dependent since SOX2^sh*Cdh1*^ daughters dispersed similar to WT clones (fig. S8B, Supplementary Video S10). Together, these data showed that E-cadherin-dependent tension differences at SOX2-WT interfaces drive preferential cohesion of SOX2 daughter cells, providing a cellular basis for clone sorting.

A direct prediction of tension-driven sorting is that at the tissue scale, overall heterotypic contact should decrease over time as clones “de-mix”. We therefore quantified the ratio of heterotypic contact perimeter to total labelled cell area, a measure that normalizes for differences in clone size and density between embryos (Fig. 4C). At baseline, SOX2 mosaics exhibited a lower heterotypic contact ratio than WT-YFP control mosaics, consistent with greater segregation from wild-type neighbors. Time-lapse imaging over twelve hours revealed that this ratio progressively decreased in SOX2 mosaics but not in YFP controls. Together these data provide direct quantitative evidence that SOX2 clones actively de-mix from their wild-type neighbors over developmental time.

Our data established a paradox: SOX2 EpSC elimination required contact with wild-type neighbors, yet the same tension differences that drove elimination also promoted clonal clustering. Thus, the more a SOX2 clone expanded, the fewer of its progeny contacted wild-type cells. We therefore predicted that elimination efficiency would decline as SOX2 EpSCs became more abundant. Increasing LV-Cre titer to elevate SOX2 representation (Fig. 5A, fig. S9A), we performed comparative growth assays against similarly transduced H2B-RFP controls. In contrast to sparse clonal results (Fig. 1B), the SOX2:WT ratio became constant when SOX2 cell density expanded across the epidermis (Fig. 5A). Per-cell SOX2 expression was comparable across these conditions, confirming that the phenotypic switch reflects neighborhood composition rather than elevated SOX2 expression (fig. S9A). Consistent with loss of loser status, SOX2 EpSCs in SOX2-dominated tissue no longer underwent significant apoptosis (Fig. 5B). Increasing homotypic SOX2 neighborhood composition was therefore sufficient to rescue loser fate.

**Figure 5.**
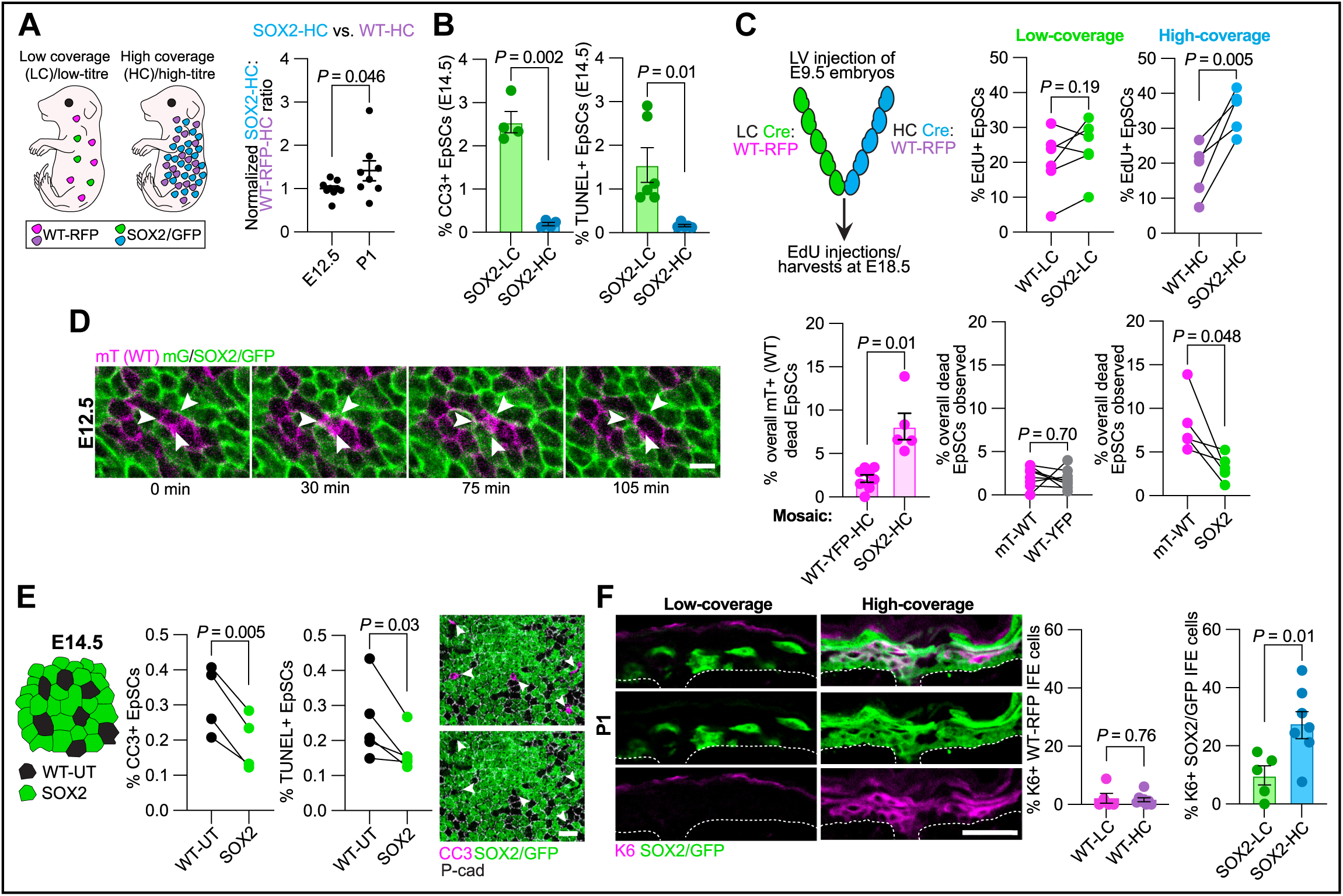
Collapse of mechanical surveillance at high SOX2 coverage enables transition to super-competition and hyperplasia. **A) Left,** schematic of low-coverage (LC) and high-coverage (HC) SOX2 EpSCs in mosaic embryos. **Right,** comparative growth assay showing that when SOX2 EpSCs are seeded in embryonic skin at high-coverage, their representation is maintained relative to similarly transduced WT controls. *n ≥* 2 embryos per litter; 2 litters per time point; two-tailed unpaired *t-*test with Welch’s correction. **B)** Quantifications of CC3 and TUNEL signals between low and high-coverage SOX2 EpSCs in E14.5 embryos. Low coverage data are re-graphed from Fig. 1D. *n ≥* 2 embryos per litter; 2 litters total; two-tailed unpaired *t*-test with Welch’s correction. **C) Left,** lentiviral transduction strategy for direct comparison of LC and HC SOX2 EpSCs. WT control EpSCs within each embryo were analyzed as controls. **Right,** quantification of EdU incorporation between SOX2 and internal, similarly-titered WT controls. *n ≥* 2 embryos per condition per litter; 2 litters total; two-tailed paired *t-*test. **D) Left,** live imaging of epidermis from mosaic E12.5 embryos containing high-coverage SOX2-eGFP EpSCs. Arrowheads denote a representative dying mT+ WT EpSC, seen when surrounded by SOX2 EpSCs. **Middle left,** quantifications of dying WT EpSCs in LV-Cre transduced at high titer into *R26-lox-stop-lox-YFP* and *R26-lox-stop-lox-Sox2-IRES-eGFP* embryos. *n ≥* 4 FOVs per embryo, 2 embryos total (WT); 1 FOV per embryo, 5 embryos total (SOX2); two-tailed unpaired *t-*test with Welch’s correction. Within-FOV comparisons of mT+ WT and WT-YFP **(middle right)** mT+WT and SOX2-eGFP death **(right)**; two-tailed paired *t*-test. **E) Left,** schematic of high-coverage SOX2 EpSCs surrounding WT-untransduced (WT-UT) EpSCs. **Middle,** quantifications reveal a marked increase in cell death when WT EpSCs are surrounded by SOX2 neighbors. *n ≥* 2 embryos per litter; 2 litters per mosaic; two-tailed paired *t-* test. **Right,** representative image of E14.5 epidermis used to collect data in **(E)**. White arrowheads denote unlabelled CC3+ WT EpSCs. **F) Left,** representative sagittal images of LC and HC SOX2 mosaic epidermis immunolabelled for K6, a marker of suprabasal hyperplasia. **Middle,** quantification of K6 expression within WT-RFP clones, verifying that K6 is not induced by LV transductions. **Right,** quantification of K6+ SOX2 clones present in low or high-coverage environments. *n ≥* 2 embryos per condition per litter; 2 litters total; two-tailed unpaired *t-*test. All data shown as mean ± s.e.m. Scale bars: 10 µm **(D)**, 20 µm **(E),** and 50 µm **(F)**.

Finally, we asked whether SOX2 dominance translates into pathological cellular behavior. To enable litter-matched comparisons, we transduced half the embryos in each litter with high-titer and the other half with low-titer LV-Cre, alongside an internal, correspondingly-titerd control LV-H2B-RFP in each embryo. In contrast to the comparable EdU incorporation seen when SOX2 EpSCs were sparse, SOX2 EpSCs in SOX2-dominated neighborhoods showed elevated EdU incorporation relative to wild-type counterparts (Fig. 5C, fig. S9B). Thus the hyperplastic potential that had been suppressed by mechanical cell competition when SOX2 EpSCs were sparse was now unleashed. SOX2 clones in SOX2-dominated tissue also displayed elevated basal-layer retention (fig. S9C), further reflecting their expanded occupancy within the EpSC compartment.

These pro-growth behaviors were also now accompanied by active elimination of WT neighbors. Live imaging of E12.5 embryos revealed a significant increase in caspase-dependent death among mTomato+ WT EpSCs in SOX2-dominated tissue compared to controls (Fig. 5D, fig. S9D, Supplementary Video S11), and CC3/TUNEL labelling of fixed epidermis confirmed that WT EpSCs were now more likely to die than their abundant SOX2 neighbors (Fig. 5E, fig. S9E). Pre-malignant SOX2 EpSCs therefore not only shed their loser status but acquired a supercompetitive winner phenotype that actively eliminated wild-type EpSCs from the tissue. This phenotype was marked by Keratin 6 (K6), a hallmark of hyperplastic epidermis, which emerged only when the SOX2 neighborhood composition was elevated (Fig. 5F, fig. S9F). Together, these data provided compelling evidence that the collapse of mechanical surveillance is not a passive loss of restraint but rather the gateway to active tumor-initiating behavior: pre-malignant EpSCs transition from loser to supercompetitor and drive hyperplastic expansion at the expense of their healthy wild-type neighbors.

## Discussion

Our findings expose E-cadherin as a master regulator of epithelial quality control in mammalian epidermis, coordinating intercellular tension at heterotypic interfaces to mark cells for competitive elimination (fig. S9). Strikingly, quantitative deviation in either direction from the neighborhood norm triggers elimination through mechanically distinct routes: increased junctional tension, associated with cell stretching upon E-cadherin excess, and reduced junctional tension and elimination upon E-cadherin suppression, the latter reminiscent of recent *in vitro* findings(*29*). While bidirectional cadherin-based elimination has precedent in the correction of morphogen-patterning noise during zebrafish development(*31, 32*), our findings in mammalian skin establish E-cadherin-dependent mechanical heterogeneity as a general and tuneable principle of pre-malignant fitness control.

Although elevated contractility and tension at boundaries between wild-type and mis-specified or pre-tumoral cells has been observed across diverse contexts and driven by a variety of upstream inputs,(*14, 26, 28, 29, 31, 49*) our findings go well beyond extending these principles to mammalian epidermis. Among our most important findings is that heterotypic contact drives redistribution of E-cadherin, thereby generating the tension heterogeneity that ultimately eliminates the less fit loser cell but without any changes in E-cadherin gene or protein expression. At present, the changes we see in cell area are only a correlate rather than a direct readout of the tension differences that track competitive fate, since instantaneous recoil reports junctional line tension, while apical area reflects the balance of junctional and medioapical cortical tension together with stress imposed by neighbors. While beyond the scope of the current study, it will be interesting to see in the future if these changes reflect volumetric stretching or apical redistribution, and how medioapical contractility is patterned in 3D at competitive interfaces.

In the basal epidermis of the skin, E-cadherin levels and interface tension correlated positively within the basal epidermis, which differed from several other model epithelia, where elevated cadherin engagement typically accompanies reduced cortical tension(*51–53*). Whether this reflects the specific context of stratifying epithelium or other distinctive physical features of this tissue, remains a provocative unknown. This said, the relevant E-cadherin mediated signal does not appear to be adhesion asymmetry per se, since inverting E-cadherin surface expression differences was not sufficient to reverse winner/loser fates between heterotypic neighbors. By contrast, relative changes in tension at heterotypic borders consistently tracked with an epidermal clone’s competitive fate.

A fascinating outcome of the mechanical framework we unearthed is that it carries an inborn vulnerability rooted in the very properties that make it effective. Because tension differences at heterotypic interfaces simultaneously mark pre-malignant cells for elimination and favour their cohesion with one another, clonal expansion inevitably works against the error correction mechanism itself. Tension-driven clustering erodes the requisite heterotypic contacts for cell competition, and the molecular and mechanical hallmarks of loser fate are progressively lost, thus facilitating the transition to super-competition.

This quality control collapse is instructive when set against other competitive systems. Pre-malignant SOX2 EpSCs minimize cell contact to escape competition, whereas in the fly notum, dMyc-driven super-competitors maximize heterotypic contact to exploit competition for expansion(*24*). Despite these distinct strategies, both converge on the same outcome, disrupted tissue homeostasis. By contrast, in the fly wing disc, failure of error correction at larger clone sizes culminate in cyst formation, retaining aberrant cells but physically walling them off from the surrounding epithelium(*28*). In mammalian epidermis, there is no such containment; instead, when heterotypic contact is lost, the spatial constraint on pre-malignant cells dissolves entirely. Group-protective effects have also been observed in the wing disc, where larger mutant patches evade elimination yet remain losers at points of wild-type contact(*54*). In comparison, SOX2 EpSCs do not merely escape elimination, they acquire a winner phenotype that actively drives apoptosis of wild-type neighbors, a more complete and pathologically consequential breakdown of tissue quality control.

While the mammalian epidermis has optimized E-cadherin-mediated tension in cell competition, it also likely becomes a double-edged sword in numerous pathological states, including wound healing, chronic inflammation, and early oncogenic transformation. The bidirectional nature of the effect we uncover predicts that any perturbation sufficient to alter E-cadherin-dependent tension at cell interfaces — whether through transcriptional, post-translational, or mechanical means — could engage competitive elimination. Taken together, our results reveal a fundamental feature of epidermal homeostasis, in which cells actively monitor the mechanical heterogeneity of their cellular constituents, deploying tension-based error correction to purge abnormal cells before they can compromise tissue integrity. Understanding how the threshold for this program is set, and how it disintegrates during pathological remodelling, may ultimately reveal new vulnerabilities at the earliest stages of tumor initiation.

## METHODS

### Mouse lines and lentiviral constructs

The following previously generated mouse lines were used in this study: *Rosa26Tomato-eGFP* (Rosa26-mTmG(*55*); Jax strain #007576), *Rosa26Brainbow2.1*(*50*) (Jax strain #013731), *Rosa26-CAG-loxP-stop-loxP-Sox2-IRES-eGFP*(*56*) (B. Hogan), *Keratin14-CreER* (Fuchs Lab-generated transgenic line), and *Rosa26eYFP*(*57*)(Jax strain #006148). For most experiments, *B6 R26^fl/fl^-Sox2-IRES-eGFP* males were bred to CD1 (CD1 ICR Charles River Laboratories) wild-type females to generate Sox2*^+/fl^* mice. B6 R26*^fl/fl^*-YFP males were bred to CD1 wild-type females for controls. Wild-type B6 males were mated to CD1 wild-type females for E-cad overexpression experiments.

All mice were housed in the Association for Assessment and Accreditation of Laboratory Animal Care International (AAALAC)-accredited Comparative Bioscience Center at the Rockefeller University. All procedures were performed using Institutional Animal Care and Use Committee (IACUC)-approved protocols and in accordance with the procedures outlined in the Guide for the Care and Use of Laboratory Animals. All mice were bred and maintained under a strict 12 hr light cycle and fed with standard chow. The temperature of the animal rooms was 20-26°C, and the humidity was 30-70%. Adult mice were housed in cages with a maximum of five mice, while LV-injected females were housed individually with handwarmers to promote viability and recovery.

Amniotic sacs were injected with lentivirus at 9.5 days post-coitum as previously described(*39*). To induce recombination of transgenic cassettes, the following lentiviruses were injected: LV-Cre, LV-H2B-RFP. Both LVs contain a shScr hairpin under U6 control. shRNA clones were obtained from the RNAi Consortium (TRC) shRNA library (Sigma), present in the pLKO.1-puro vector. The puro cassette was swapped out for an H2B-RFP marker before transfection into low-passage HEK-293-FT cells for high-titer lentivirus production which could then be transduced in vivo. Knockdown efficiency of *Cdh1* was confirmed in both low-passage keratinocytes (**fig. S4A**) and E14.5 embryos(*46*) (**fig. S4B**). To generate LV-Ecad-OE (**fig. S5A**), the coding sequencing of mouse Cdh1 from a previously validated E-cad-tdTomato fusion protein construct(*48*) (Addgene #101278) into an LV-GFP construct to generate an E-cad-EGFP fusion protein expressed under PGK control.

### Tamoxifen and EdU administration

For the postnatal experiments performed in **fig. S1A**, the backskins of P49 *K14CreER;R26-lox-stop-lox-Sox2-IRES-eGFP* mice were shaved and treated topically with 50 µg/g body weight tamoxifen (Sigma Aldrich) in 100% ethanol daily for 5 days. Three weeks post-tamoxifen treatment, mice were sacrificed and backskins harvested. For lineage tracing experiments, 200 µL of tamoxifen dissolved at 2 mg/mL in corn oil was intragastrically administered to pregnant females at embryonic day E14.5. For labelling of proliferating cells in embryonic epidermis, EdU (25 µg/g; ThermoFisher) was administered via intraperitoneal (i.p.) injection of pregnant females. Dosages were split between both i.p. cavities to ensure equivalent administration to both uterine horns. 30 min later, animals were sacrificed according to AAALAC guidelines and embryos were dissected from their uterine horns. Backskins were then processed.

### Drug treatments

Dynasore (Abcam; 250 µM) and Z-DEVD-FMK (Tocris; 20 µM) were dissolved in agarose-medium solution (see live imaging) and allowed to incubate for 3 hr prior to fixation or live imaging. 0.5% DMSO was used as a control for both treatments. For fixed-tissue characterization of apoptosis at E12.5 with DEVD, embryos were incubated for 15 hr.

### shRNA sequences

*Cdh1* shRNA: 5’-CCGAGAGAGTTACCCTACATA-3’ (as used in Lay et al(*46*)). Scramble shRNA 5’-CAACAAGATGAAGAGCACCAA-3’ (as used in Beronja et al(*39*)).

### Immunofluorescence and antibodies

For all immunofluorescence experiments, embryos were fixed for 1 hr in 4% paraformaldehyde (PFA) at room temperature (RT). Adult backskin biopsies were fixed for 30 min in 4% PFA at 4°C. For whole-mount of imaging of backskin at all time points except E12.5, backskin was dissected from the animal and cut into four quadrants. Whole-mounts of E12.5 embryos were kept intact throughout staining and imaging. After fixation, samples were permeabilized in 0.3% PBS-Triton (PBS-T) for 1 hr at RT and blocked in blocking buffer (5% donkey serum, 2.5% fish gelatin, 1% BSA, 0.3% PBS-T) for 30 min. Primary antibodies were incubated overnight at 4°C and washed for ≥ 20 min in PBS-T. Secondary antibodies were incubated overnight at 4°C and washed for ≥ 20 min in PBS-T. Whole embryos and backskins were mounted in Prolong Diamond Antifade Mountant (Invitrogen) for imaging. For sections at P1, backskin was cut into strips, embedded and frozen in OCT (Leica), and sectioned with a Leica cryostat (12 µm sections). EdU labelling of embryos was performed using the Click-iT Alexa Fluor 647 Imaging kit (ThermoFisher) between application of the primary and secondary antibodies. Our protocol followed manufacturer’s instructions except that we incubated the EdU detector for 2 hr instead of 45 min. For TUNEL labelling, the Click-iT TUNEL Alexa Fluor 647 Imaging Assay for Microscopy and HCS kit (Invitrogen) was used and followed according to manufacturer’s instructions. Antibodies were used as follows: rat anti-RFP (Chromotek, 1:500), rabbit anti-RFP (MBL, 1:500), goat anti-RFP (MyBioSource, 1:200), chicken anti-GFP (Abcam, 1:1,000), rat anti-E-cadherin (M. Takeichi, 1:500), rabbit anti-keratin10 (Covance, 1:500), rabbit anti-cleaved-caspase3 (Cell Signaling, 1:1,000), goat anti-P-cadherin (R&D, 1:200), rat anti-ITGA6 (BD Pharmingen, 1:1,000), and rabbit anti-Keratin6A (BioLegend, 1:500). All secondary antibodies were raised in a donkey host and conjugated to DyLight 405 Anti-Rabbit (Jackson ImmunoResearch, 1:100), AlexaFluor488, AlexaFluorRRX, or AlexaFluor647 (Jackson ImmunoResearch, 1:500). Rhodamine phalloidin and Alexa Fluor 647 (both Life Technologies, 1:100) was used to label F-actin. DAPI was used to label nuclei (1:5,000).

### Microscopy and live imaging

E12.5 live imaging was carried out according to Ouspenskaia et al(*58*). Embryos were placed on their sides in a 35-mm Lumox-bottom dish (Sarstedt). Embryos were immobilized in a custom-built mold and stabilized in an agarose solution composed of 2% low-melting SeaPlaque Agarose (Cambrex) in a solution of epidermal culture medium supplemented with 300 µM calcium chloride. After equilibrating at 37°C and 5% CO_2_ for 3 hr, imaging was performed using an inverted spinning disk confocal system (Andor Dragonfly 202) at 15-minute intervals for 12 hr (488 and 561 nm laser beams, 20X air objective, NA = 0.75). Time-lapse images were acquired with a Zyla cCMOS camera (Andor). Approximately 3-5 non-overlapping regions were filmed from each embryo. Embryos were maintained at 37°C and 5% CO_2_ throughout the experiment.

All whole-mount and epidermal cryosections immunofluorescence images were acquired using the same spinning disk confocal system as described previously and a 20X air (NA = 0.75), 40X air (NA = 0.85), or 63X oil objective (NA = 1.4). All images were assembled and processed using ImageJ and/or Imaris.

### Fluorescence Activated Cell Sorting and RNA extractions

Dissociation of E14.5 backskins was performed using a solution of 0.25% Trypsin/Versene (1:1; Invitrogen) incubated for 20 minutes at 37°C. Following filtration, single cells were stained with the following: CD45-PE/Cy7 (BioLegend, 1:200), CD117-PE/Cy7 (BioLegend, 1:200), CD31-PE/Cy7 (BioLegend, 1:200), CD140a-PE/Cy7 (Invitrogen, 1:200), DAPI (1:10,000), CD49f-PerCP/Cy5 (BioLegend, 1:1000), and CD29-APC/Cy7 (BioLegend, 1:1000) for 20 minutes on ice. Cells were then sorted using a BD FACSAriaII and/or SONY MA900 directly into RT Tri-Reagent (Zymo Research), flash-frozen, and stored at -80°C. Two biological replicates were used per condition, with at least two embryos pooled per biological replicate. RNA extractions were performed on purified EpSCs (α6+β1+XFP+) and total RNA was isolated with a DirectZol RNA MiniPrep Kit (Zymo Research) as per manufacturer’s instructions.

### Statistics and reproducibility

Experiments were repeated using at least two independent litters (≥ 2 animals per litter) per experiment. Statistical methods were not used to predetermine sample size. Statistical and graphical analyses were performed in GraphPad Prism (9/10). All datasets generated were tested for normal distribution using Prism 9/10 (GraphPad) and all data sets that failed this test were subject to non-parametric tests for further analysis. Analysis of all live imaging and laser ablation data was performed with investigators blinded to genotype. Importantly, the lentiviral injection specialist was blinded to experiment allocation and results. All other experiments were not randomized and the investigators were not blinded to allocation nor outcome assessment, given the lack of ambiguity in phenotypes observed and internal controls used. Sample sizes, replicates, and statistical tests used are indicated in each figure legend. All experiments reported were replicated at least twice with uniformly reproducible results.

### Image processing and analysis

#### Adult hyperplasias

Maximum projection images were assembled from cryosection images of second-telogen treated with tamoxifen in ImageJ. Hyperplasias were defined as continuous segments of epidermis with multi-layered (more than one) α6+ staining. Overgrowths and GFP fractions were manually traced as a fraction of total α6+ interfollicular epidermis.

#### Comparative growth assays

Maximum projection images were assembled from whole-mount images of embryos (E12.5) and sagittal cryosection images of neonatal pups (P1) infected with identical mixtures of LV-Cre and LV-H2B-RFP. Images were segmented in ImageJ to identify and count the number of GFP+ and RFP+ nuclei. GFP+RFP+ double positive nuclei were excluded. To obtain the ratios shown in **Fig. 1B**, **3E**, **5A**, and **S5F**, the ratios of GFP+ to RFP+ EpSCs were calculated at E12.5 and P1. The ratio at P1 was then normalized to that at E12.5 to determine whether the representation of GFP+ changed relative to RFP over developmental time.

#### Cell proliferation/CC3/TUNEL assays

Proliferation was measured by incorporation of labelled nucleotide analogues following a 30 min EdU pulse. Maximum projection images featuring only the basal plane of the epidermis were assembled in ImageJ. The GFP and RFP signals were thresholded to allow unambiguous detection of competing clones. CC3, TUNEL, and EdU signals were similarly thresholded. To assess differences in proliferation, measurements were paired and the incorporation of EdU by Cre+GFP+ EpSCs was directly compared to that by their Cre-RFP+ counterparts in the same embryo. Absolute EdU incorporation percentages are never compared across embryos because subtle differences in developmental timing between animals, even within the same litter. The total number of GFP+ and RFP+ TUNEL+/CC3+ bodies were then plotted as a fraction of total GFP+ or RFP+ EpSCs. GFP+RFP+ double positive nuclei were excluded. Where possible, paired comparisons were made within the same embryo to account for baseline variability in death observed between embryos.

#### Live imaging of death events

Death events were manually scored and two independent investigators were both blinded to the conditions and analyzed the data. Cells that looked unhealthy at the start of each video, folds, and areas of global death were excluded from analysis. For each putative cell death event, the entire z-stack acquired (150 µm) was scanned through to rule out extrusion of EpSCs into other planes. At the start of each movie, all GFP+ EpSCs were binned into one of three categories depending on their neighborhood conformation (**Fig. 1G**). Dying GFP+ EpSCs over the course of the movies were correspondingly categorized and reported as a percentage of all GFP+ EpSCs categorized into the original bins. Cell contact was defined as at least one shared vertex. Analysis of dying mT+WT EpSCs was performed similarly. Since understanding differences how neighborhoods change death rates is fundamental to our analysis, statistical comparisons were made on the level of field-of-view comparison, and not embryos, due to considerable variation in neighborhood composition between individual fields of view, even within the same embryo. These differences would be obscured by averaging across the whole embryo (see also superplotted data in **fig. S1E and G**).

#### Live imaging of clonal dispersion

For WT-YFP and competitive SOX2 mitotic clones, only EpSCs completely surrounded by mT+ WT neighbors were analyzed to exclude the possibility of interference from homotypic neighbor contacts. For non-competitive SOX2 mitotic clones, only SOX2 clones completely surrounded by SOX2 EpSCs were analyzed; any clones contacting mT+ WT EpSCs were excluded. Mitotic events were classified as “splits” if contiguous membrane-GFP signal encapsulated each daughter cell and if there was a visible, measurable distance separating the borders of the resulting daughter EpSCs within 45 min post-division. This was empirically determined based on comprehensive observation of endogenous dividing WT clone behavior. Distances between resulting sisters were measured manually per 15-min frame in Imaris. Data were then fit using the straight-line equation parameter of the nonlinear regression analysis function in GraphPad Prism with least squares regression without weighting. To determine whether the slope of at least one datasets differed between the three conditions, data were compared using extra sum of squares F-test. For head-to-head comparisons of conditions, data were replotted in GraphPad for each of the three comparisons (WT-YFP vs. competitive SOX2, WT-YFP vs. non-competitive SOX2, and competitive SOX2 vs. non-competitive SOX2), and the above regression analyses were rerun to determine individual *p* values. To correct for multiple comparisons, the Bonferroni correction was applied, which changed the threshold for significance (alpha) from 0.05 to 0.017. For centroid analysis in ImageJ, daughter cells were drawn as regions of interest. The (x,y) coordinates were calculated for both daughter cells and tracked over time. Distances between the resulting coordinates was then calculated over time.

#### Clone shape prediction modelling

To quantify clone tension, we performed laser ablation at three junction types: host-host (γ), clone-host boundary (γ_b), and clone-clone (γ_c) contacts, using instantaneous retraction velocity as a proxy for junctional tension. Clone tension σ was calculated according to Bosveld et al. as σ = 1/γ · {γ_b − [(γ + γ_c)/2]}, and mean tension ratios (γ_b/γ and γ_c/γ) were plotted on the phase diagram described therein to predict clone behavior. A positive σ predicts clone rounding, while a negative σ predicts clone scattering relative to the surrounding tissue.

#### Image segmentation and cellular morphometric analysis

For each z-stack, the focal plane with the highest proportion of basal cells was selected for downstream analysis. Cell segmentation was performed on this plane using a custom-trained model within Cellpose-SAM (version 4.0.6; Pachitariu et al., 2025 ((*59*)), followed by manual correction to resolve segmentation errors. The resulting segmentation masks were converted into cell outlines using a custom-written Python script. These outlines were subsequently imported into Tissue Analyzer (Fiji plugin(*60*)), which was used to extract cell- and junction-level morphological features. All extracted parameters were combined with fluorescent marker expression data in a custom-written Python script, and cells were assigned to subpopulations based on marker identity. Islands were manually identified within subpopulations. For each image, per-category means, standard deviations, and cell/junction counts were extracted. The image was treated as the unit of replication throughout. Because the number of islands per image was often small, to assess the consistency of directional effects in size difference between islands and the other two cell populations, sign tests were performed on per-image means, testing whether the proportion of images in which one category exceeded another differed from 0.5 (one-sample binomial test, two-tailed). However, the more numerous cell population values were compared using paired t-tests with Bonferroni corrections. Statistical analyses were performed in R, and plots were generated using ggplot.

#### Clustering index/clonal dispersion assay

For all clonal analyses (including Brainbow validation), lentiviruses were substantially diluted so MOI was significantly lower than 1. Clones were thus defined as a group of labelled EpSCs of a single color lying far away from other labelled clones (limited to ∼ 1 clone per 20X FOV). Moreover, to ensure growth- and developmental stage-matched comparisons, SOX2 and wild-type control clones were induced and compared within the same embryos. To describe the cohesiveness of different EpSC clones, we developed a cluster index *I*. The cluster index *I* is defined as:

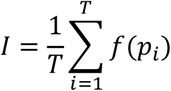

Where *T* is the total number of cells in the clone, *p_i_* is the number of directly adjacent neighbors of cell *i*, and *f*(*p_i_*) is a weighting function that assigns a value between 0 and 1 to the cell *i* depending on the number of its neighbors *p_i_*. The function *f*(*p_i_*) is defined as:

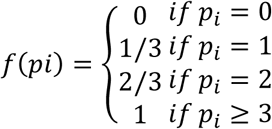

This reflects the assumption that a higher number of neighboring cells indicates a greater degree of local clustering. The cluster index *I* therefore describes the overall cohesiveness of a clone, ranging from: *I* = 0 for a fully dispersed clone (no cell has any direct neighbors), to *I* = 1 for a maximally clustered clone (every cell has at least three neighbors) (**fig. S6B**).

#### Homo/heterotypic border and intracellular E-cadherin and F-actin analysis

For *in vivo* analyses, 2-5 63X images were randomly taken per embryo and maximum projection images were assembled. 20-30 EpSCs were analyzed per embryo. For homotypic vs. heterotypic analyses, the mean gray value of the E-cadherin (or Phallodin for F-actin) signal of every border was measured. The average homotypic and heterotypic border values were calculated per cell. All values were normalized to the average WT homotypic value across the embryos across all litters. For island analyses, the mean gray value of the signal of all borders of analyzed cells was extracted as a single region of interest. All values were normalized to the average WT homotypic island value across all embryos across all litters.

#### Suprabasal hyperplasia/neighborhood analysis

For the quantification of K6+ hyperplasias, K6+ WT-RFP or SOX2-eGFP EpSCs were manually counted as a fraction of total WT-RFP or SOX2-eGFP clones in the interfollicular epidermis, irrespective of suprabasal or basal localization/identity. Neighborhoods were defined as continuous, contacting basal and/or suprabasal clones. Hair follicles were excluded from analyses. WT-RFP cells directly in contact with SOX2-eGFP clones were excluded from analyses, as were GFP+RFP+ double positive nuclei.

### Laser ablation

Laser ablation experiments were performed as described previously(*61, 62*). E14.5 embryos were prepared similarly to live imaging experiments and imaged using an inverted Zeiss LSM 880 laser scanning confocal and multiphoton microscope system. Severing of junctions was achieved using a tunable Ti-:sapphire near-infrared laser (Chameleon Ultra II, Coherent Scientific) tuned to 800 nm. Laser power was used at 100% transmission at a scan speed of 10 and pixel dwell time of 0.64 µsec. The junctional response following ablation was imaged using a LD LCI Plan-Apochromat 40X/NA=1.2 water immersion objective with correction collar the maximum possible frame acquisition rate, for up 15 frames post-ablation and at 12-bit depth. The displacement of the neighboring tricellular junctions relative to their initial position at the time of ablation was then manually tracked using Imaris. Quantification of effects of ablation was performed by two independent and blinded investigators. Data was then fit to a Kelvin-Voigt model(*42*), allowing us to extract force:elasticity ratios and tau for each class of junction. Ablations that inadvertently wounded EpSCs or failed to completely sever cell-cell junctions were excluded from analyses. No more than 12 cuts were performed per embryo and all cuts were performed on cell-cell junctions lying parallel to the mediolateral body axis. Junctions were selected for ablation across the entire back-skin, while being careful to avoid limb, head, or neck regions.

### Quantitative real-time PCR (qPCR)

#### RNA isolation

RNA was isolated from sorted EpSCs stored in Trizol at the recommended 1:3 ratio at -80°C until use. The total number of sorted cells ranged from 5000 – 50,000 EpSCs per samples. Where applicable, samples of the same genotype and lentivirus titer were pooled to reach at least 5,000 EpSCs. RNA was isolated using the Direct-zol RNA MicroPrep Kit (Zymo Research) according to manufacturer’s instructions including a 15 min treatment with DNase I. The final elution volume was 10 µL and RNA quantity and quality were measured on a NanoDrop 8000 spectrophotometer (Thermo Fisher). The same amount of RNA (∼50 ng) was loaded per sample for subsequent cDNA synthesis.

#### cDNA synthesis

∼50 ng of RNA were loaded per sample, and cDNA was synthesized using the SuperScript^TM^ VILO^TM^ cDNA Synthesis Kit (Thermo Fisher). In a final volume of 20 µL we mixed VILO Reaction mix (5X), SuperScript Enzyme (10X), RNA template and DEPC-treated water (Thermo Fisher). cDNA was synthesized using the following PCR program: 10 min at 25°C – 60 min at 42°C – 5 min at 85°C – 12°C. For all quantitative real-time PCR (qPCR) reactions cDNA was diluted to get a final concentration of 1.5 ng/µL.

#### qPCR

Equal amounts of cDNA template were mixed with validated gene-specific primers (in a final 1:100 dilution of a 10 µM solution), DEPC-treated water and Power SYBR green PCR Master Mix (2X, Thermo Fisher) in a final volume of 10 µL per sample. The amplification efficiency of every primer pair was validated to be between 90 – 110% (as tested by standard curves) in advance. qPCR was performed in technical duplicates using an Applied Biosystems 7900HT Fast Real-Time PCR machine and relative expression (RE) to a control sample was calculated with the RelQuant software (Bio-Rad Laboratories) by normalizing to *Ppib2* housekeeping gene expression.

#### qPCR Primers

*Ppib2* forward: 5’-GTGAGCGCTTCCCAGATGAGA-3’; *Ppib2* reverse: 5’- TGCCGGAGTCGACAATGATG-3’; *SOX2* forward: 5’- AACGGCAGCTACAGCATGATGC-3’; *SOX2* reverse: 5’-CGAGCTGGTCATGGAGTTGTAC-3’; *Cdh1* forward: 5’- GGGTCATCAGTGTGCTCACCTCT-3’; *Cdh1* reverse: 5’- GCTGTTGTGCTCAAGCCTTCAC GCTGTTGTGCTCAAGCCTTCAC-3’.

### Immunoblotting (western blot) and analysis

To isolate lysates for immunoblots from *in vitro* cell cultures, culture media was aspirated, wells were washed with PBS, and cells were lysed in chilled RIPA buffer containing cOmplete protease inhibitors and PhoSTOP phosphatase inhibitors (Roche). Protein was quantified using a Pierce™ BCA protein quantification kit and BioTek CyTation (Agilent). 20 μg protein was loaded for SDS– PAGE and run in MOPS buffer. To isolate cells directly from the epidermis of live E14.5 embryos for immunoblotting, WT and SOX2+ EpSCs were purified as above and sorted into cold FACS buffer, washed with cold PBS, and centrifuged at 300 x *g* for 5 min. Cells were resuspended at a concentration of 1000-2000 cells/µL and lysed as above. 10,000 cells were loaded per well for SDS-Page and run in MOPS buffer. The following primary antibodies and dilutions were used for *in vitro* experiments: E-CAD (rabbit, 1:1,000, Cell Signaling); SOX2 (rabbit, Abcam, 1:2000), eIF2*α* (rabbit, 1:5,000, Cell Signaling), PSAT1 (mouse, 1:500, Abnova), and H3 (mouse, 1:5000, Cell Signaling). LiCOR secondary antibodies IRDye 800CW donkey anti-rabbit IgG and IRDye 680 LT donkey anti-mouse IgG were used at 1:10,000.

Wet transfer was performed overnight at 4°C at 30V. PVDF membranes were blocked in LiCOR blocking buffer. Membranes were incubated in primary antibody overnight in blocking buffer with 0.2% Tween-20 at 4 °C, washed in PBS with 0.2% Tween-20, and incubated in secondary antibody diluted in blocking buffer with 0.2% Tween-20 and 0.01% SDS for 1 hr at room temperature. The following primary antibodies and dilutions were used for *in vivo* experiments: ECAD (rabbit, 1:500, Cell Signaling); SOX2 (rabbit, Abcam, 1:1000), SDHA (mouse, Abcam, 1:1000), and H3 (mouse, 1:1000, Cell Signaling). LiCOR secondary antibodies IRDye 800CW donkey anti-rabbit IgG and IRDye 680 LT donkey anti-mouse IgG were used at 1:5,000.

Membranes were imaged via fluorescence on an Odyssey CLx Imager and processed using Adobe Photoshop CS5. Bands were quantified semi-quantitatively using densitometry in Fiji.

### Statistical modelling of SOX2-EGFP and WT-RFP epidermal progenitor dynamics

To model the relative growth of SOX2-EGFP and WT-RFP subpopulations during epidermal development, we constructed a two-phase discrete-time population model informed by empirically measured cell death rates and published tissue growth parameters. Both labelled populations were initialized at 10% of total epidermal progenitors at E12.5, with the remaining 80% comprising unlabelled cells. The model was parameterized using the two-phase framework described by Damen *et al*(*63*), in which the net tissue growth rate (λ) was measured empirically as λ₁ = 0.66/day during the early amplification phase (E12.5–E15.5) and λ₂ = 0.70/day during the subsequent expansion phase (E15.5–P1, where E19.5 corresponds to P0). Critically, λ describes net tissue expansion rather than raw proliferation. The underlying proliferation rate was therefore back-calculated as λ plus the empirical WT-RFP death rate, yielding an implied proliferation rate of 0.699/day in Phase 1 and 0.70/day in Phase 2. All populations were assumed to share the same proliferation rate; differences in net growth arise solely from differences in death or differentiation rates. The model was run in discrete 12 hr timesteps.

Death rates for WT-RFP and SOX2-EGFP cells were derived from time-lapse imaging experiments in which dying cells were counted within a field of view over 12-hour movies and expressed as a percentage of the starting visible population. Mean death rates were 1.97%/12h (95% confidence intervals (CI): 1.28–2.66%; co-efficient of variation (CV) = 88%, n = 20 movies from 5 embryos) for WT (RFP) and 5.4%/12h (95% CI: 3.40–7.40%, CV = 75%, n = 24 movies from 9 embryos) for SOX2 (EGFP). See also superplotted data in **fig. S1E**. In Phase 2, cell death was set to zero in all populations, consistent with the cessation of apoptosis after E15.5. Basal progenitors committed to differentiation with probability r = 1/3 per division cycle (also based on Damen(*63*)), and an additional SOX2-EGFP-specific differentiation rate was estimated by fitting the model to the experimentally measured SOX2-EGFP:WT-RFP ratio at P1 (*n* = 7 animals) using bounded scalar optimisation (scipy.optimize.minimize_scalar). The best-fit extra differentiation rate was 13.82%/day (95% CI: 11.16–16.38%/day, where the CI was propagated from the 95% CI of the empirical death rates; see **fig. S2E**). To decompose the relative contributions of Phase 1 death and Phase 2 differentiation to the total ratio drop, simulations were run in which each perturbation was removed independently. All modelling and statistical analyses were performed in Python 3 using NumPy, SciPy, and Matplotlib.

### Artificial Intelligence Tools

During the preparation of this manuscript, the authors used Claude (Anthropic, claude-sonnet-4-6) to assist with manuscript writing, including drafting and editing text for clarity and concision, and to generate Python and R codes used in the statistical analysis, graphic generation, stochastic population dynamics modelling (**fig. S2**) and phase diagram analysis **(fig. S6A**). All AI-assisted text was reviewed and substantially revised by the authors, all quantitative outputs and model results were independently verified, and the authors take full responsibility for the accuracy and integrity of the final published work.

## AUTHOR CONTRIBUTIONS

E.A.N.T., S.J.E., and E.F. conceived the experiments and wrote the manuscript. C.S. assisted with all live imaging and laser ablation analyses, as well as additional image analysis. T.O. performed all lentiviral injections. M.P., S.M., M.S., C.J.B., and A.G. assisted with image analysis. A.R.B. and S.M.P. assisted with flow cytometry experiments. A.R.B. assisted with molecular cloning. K.S.S. and S.M.S. provided conceptual input and additional experimental support. M.S. performed qPCRs. J.S.S.N. assisted with Western blots, statistical analyses and tissue harvests. P.B. assisted with laser ablation experiments and video processing. E.A.N.T. and S.J.E. performed all remaining experiments, data analyses, and quantifications. All co-authors provided input on the manuscript.

## MATERIALS AND CORRESPONDENCE

Correspondence to Stephanie J. Ellis or Elaine Fuchs

## ACKNOWLEDGEMENTS

We thank A. North and M. Alfonso of the Rockefeller University (RU) Bio-Imaging Resource Center for assistance with laser ablation experiments (RRID:SCR_017791); L. Polak and L. Hidalgo for animal assistance; RU Comparative Bioscience Center (AAALAC-accredited) for care of mice in accordance with National Institutes of Health (NIH) guidelines; L.B. Vosshall, A.E. Shyer, P. Cohen, D.L. Moore, A. Sharma, K.M. Plachta, R.L. Kimura, Y. Chiba, P. Thompson, D.L. Wang, and all members of the Fuchs and Ellis labs, specifically R. Niec, L. Polak, A. Khandekar, S. Sharma, S. Yuan, J. Almagro, M.D. Abdusselamoglu, C. Xu, W. Song, and M. Kudelka for discussion and/or input on the manuscript. E.A.N.T., S.M.S., and C.J.B. are supported by NIH Predoctoral Individual National Research Service Awards (NRSA) (F31-AR085502, F31-AR083275, and AR087287 respectively). C.S. is the recipient of a Max Perutz PhD Fellowship (Universität Wien). K.S.S. received support from a NYSCF Druckenmiller Postdoctoral Fellowship. M.S. was supported by a RU Women & Science Graduate Fellowship and a Boehringer-Ingelheim Fonds PhD Fellowship. J.S.S.N. was supported by a NIH Predoctoral Individual NRSA (F30-HD107964) and T32-GM007739. S.M.P. is supported by a NIH Pathway to Independence Award (K99-AR084573). S.J.E. is a Vallee Foundation Scholar. Research in the S.J.E. lab was funded in part by the Austrian Science Fund (FWF) [grant DOI: 10.55776/STA54]. S.J.E. received support from a NYSCF Druckenmiller Postdoctoral Fellowship and a NIH Pathway to Independence Award (K99-AR074557). E.F. is an HHMI Investigator. This research was supported by the NIH (R37-AR27883 to E.F.).

## SUPPLEMENTAL FIGURES LEGENDS

**Supplemental Figure 1.**
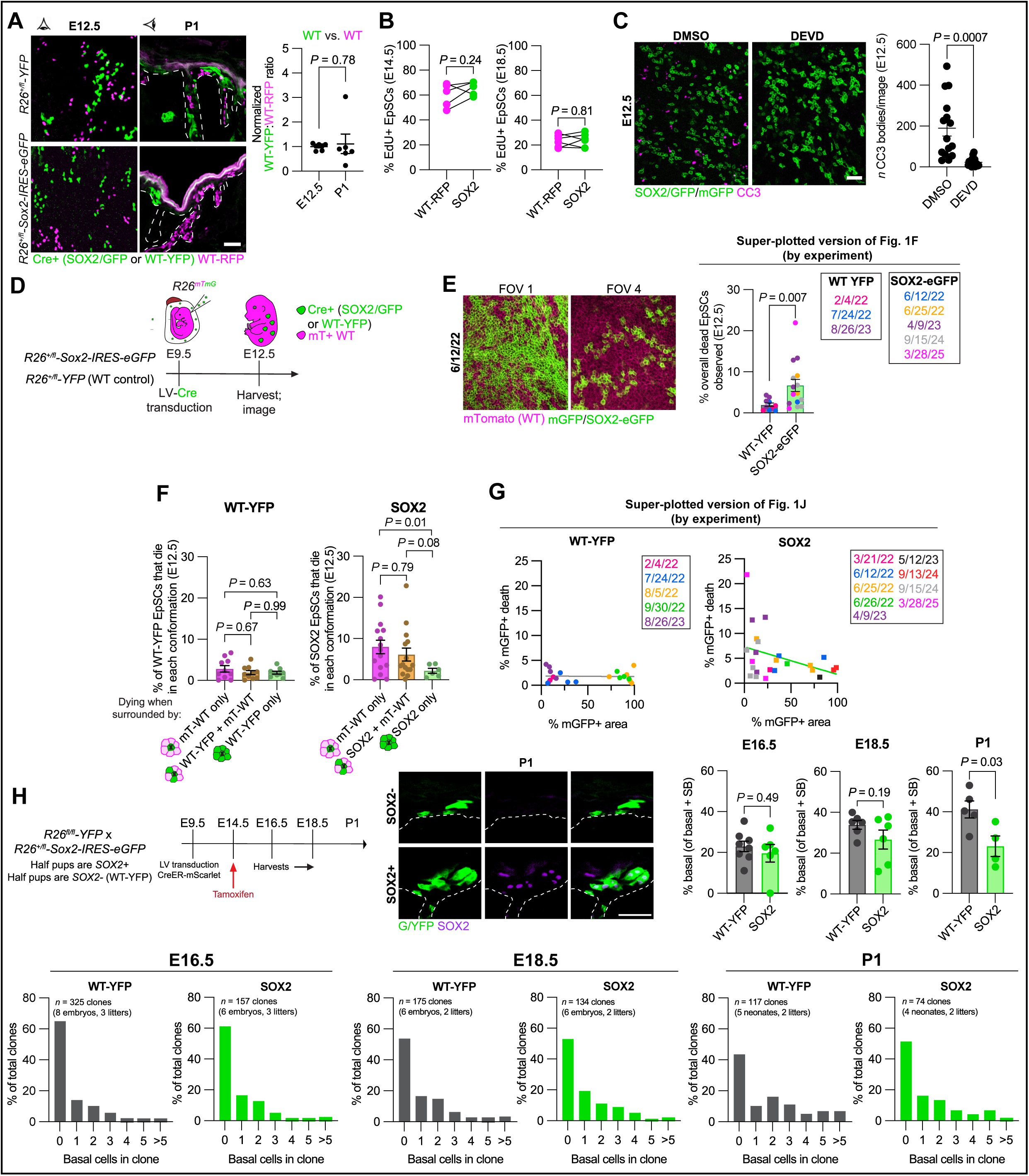
Validation of SOX2 EpSC fitness deficit, caspase-dependence, and clonal behavior across epidermal development. **A) Left,** representative immunofluorescence images of CGAs in *R26-lox*-*stop-lox-YFP* animals **(top row)** comparing WT-YFP vs. WT-RFP EpSC fitness and accompanying quantification **(right)**. *n ≥* 2 embryos per litter; 2 litters per time point; two-tailed unpaired *t-*test with Welch’s correction. **Bottom row,** representative images of SOX2 comparative growth assay performed in **Fig. 1B**. Dashed lines denote epidermis-dermis border. **B)** EdU incorporation assays comparing WT and SOX2 EpSC proliferation at E14.5 **(left)** and E18.5 **(right)**. *n* ≥ 2 embryos per litter; 2 litters per time point; two-tailed paired *t*-test. **C) Left,** representative whole-mount maximum projection immunofluorescence images of CC3-stained epidermis of DMSO or DEVD-treated E12.5 embryos. **Right,** quantification of CC3+ bodies per image. *n =* 1-4 images per embryo, 2-3 embryos per litter, 2 litters per condition; two-tailed unpaired *t-*test with Welch’s correction. **D)** Schematic of E12.5 live imaging experimental methodology. **E) Left,** two representative, differentially transduced 20X FOVs taken from a single embryo, demonstrating that individual FOVs from the same embryo are not technical duplicates given the disparate neighborhood conformations that arise from lentiviral transduction. **Right,** superplot of **Fig. 1F** where the FOVs from each individual experiment (embryo) are color-coded. **F)** Comparison of conformation-dependent death in E12.5 WT-YFP **(left)** and SOX2 **(right)** EpSCs (re-graphed from **Fig. 1, F and J**). **G)** Superplotted versions of **Fig. 1J** separated by genotype (WT-YFP on **left;** SOX2 on **right**) where each experiment (embryo) and its corresponding FOVs are color-coded. **H) (Top row) Left,** lineage tracing experimental strategy to compare differentiation propensities of litter-matched SOX2 vs. WT-YFP control EpSCs. **Middle,** representative sagittal immunofluorescence images of clones from SOX2-**(top)** and SOX2+ **(bottom)** littermates at P1. Dashed lines indicate the epidermis-dermis border. **Right,** quantification of basal EpSC population relative to total basal and suprabasal population of labelled clones at E16.5, E18.5, and P1. *n* ≥ 2 embryos per litter, 2 litters per time point; two-tailed unpaired *t-*test with Welch’s correction. **Bottom row,** distribution frequencies of clone size by indicated time point. 0 basal cells indicates that all clonal progeny were suprabasally-localized. Sample sizes are indicated on graphs. All data shown as mean ± s.e.m. Scale bars = 50 µm.

**Supplemental Figure 2.**
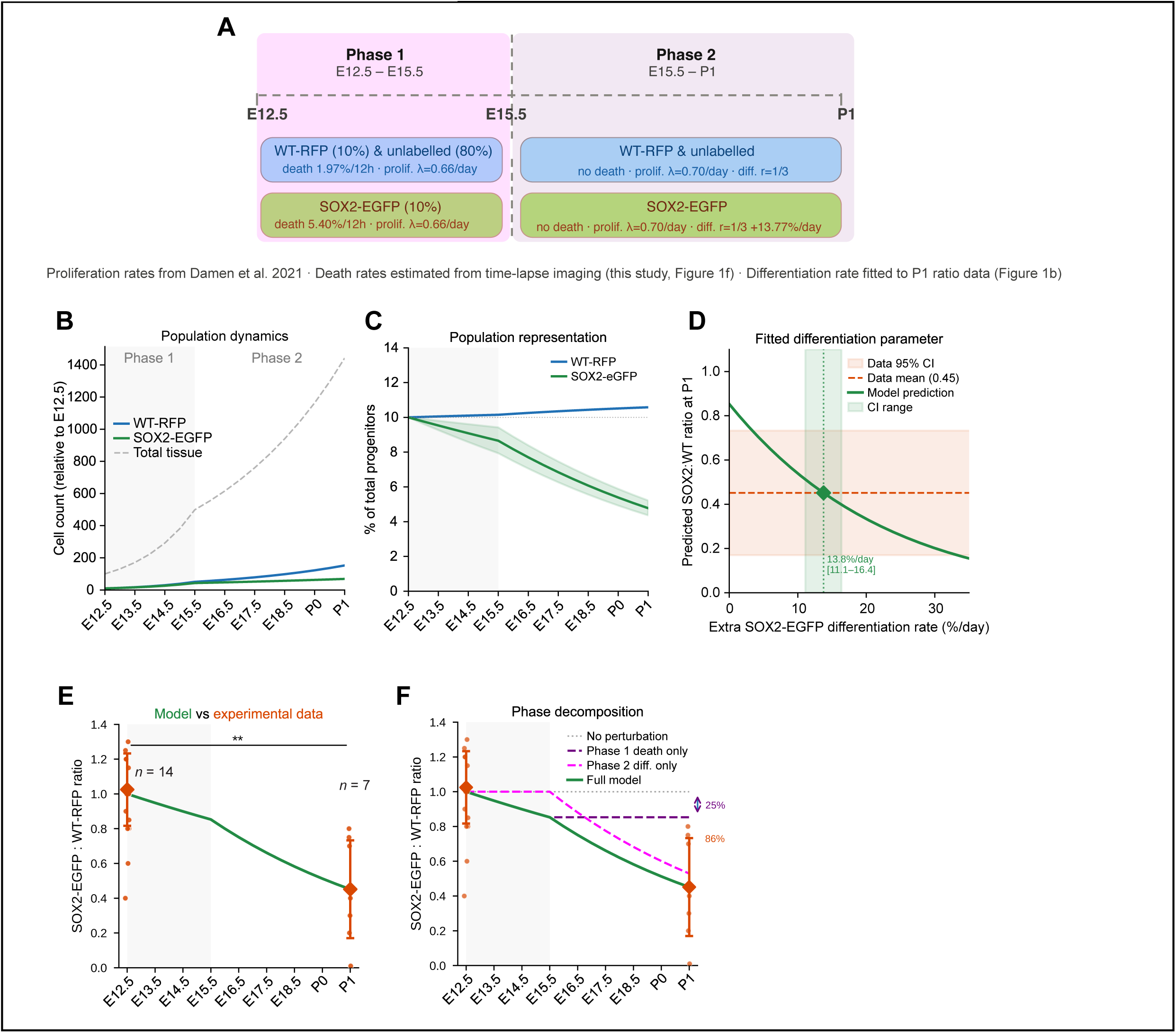
Statistical modelling of SOX2 EpSC population dynamics across epidermal development. **A)** Model schematic. Phase 1 (E12.5–E15.5): WT-RFP (10% labelled) and unlabelled WT EpSCs (80%) die at 1.97%/12hr while SOX2 EpSCs (10% labelled) die at 5.40%/12hr; all populations proliferate at λ = 0.66/day. Phase 2 (E15.5–P1): apoptosis is negligible and all populations proliferate at λ = 0.70/day; baseline differentiation occurs at r = 1/3, with SOX2 EpSCs differentiating at an additional 13.77%/day. Proliferation rates derived from Damen et al. 2021; apoptosis rates from time-lapse imaging (**Fig. 1F**); differentiation rate fitted to P1 ratio data in **Fig. 1B**. **B)** Modeled population dynamics across development. Cell counts shown relative to E12.5; total tissue (dashed grey), WT-RFP (blue), SOX2-eGFP (green). **C)** Modeled population representation as percentage of total progenitors over time. Shaded regions indicate model variance. **D)** Fitted Phase 2 differentiation parameter. The predicted SOX2 : WT ratio at P1 is shown as a function of the additional SOX2-eGFP differentiation rate. Orange dashed line indicates the experimentally observed mean P1 ratio (0.45); shaded region shows 95% CI. The fitted parameter (13.77%/day, 95% CI: 11.1–16.4) is indicated by the green dotted line. **E)** Comparison of modelled SOX2/GFP:WT-RFP ratio (green line) to experimentally measured ratios at E12.5 (*n* = 14) and P1 (*n* = 7). Orange diamonds indicate mean experimental values; error bars show SD; Mann-Whitney U-test. **F)** Phase decomposition. Modeled SOX2:WT ratio trajectory under three scenarios: no perturbation (grey dotted), Phase 1 apoptosis alone (purple dashed), Phase 2 differentiation alone (pink dashed), and the full model incorporating both (green solid). Percentages indicate the fraction of full-model elimination achieved when each phase operates in isolation.

**Supplemental Figure 3.**
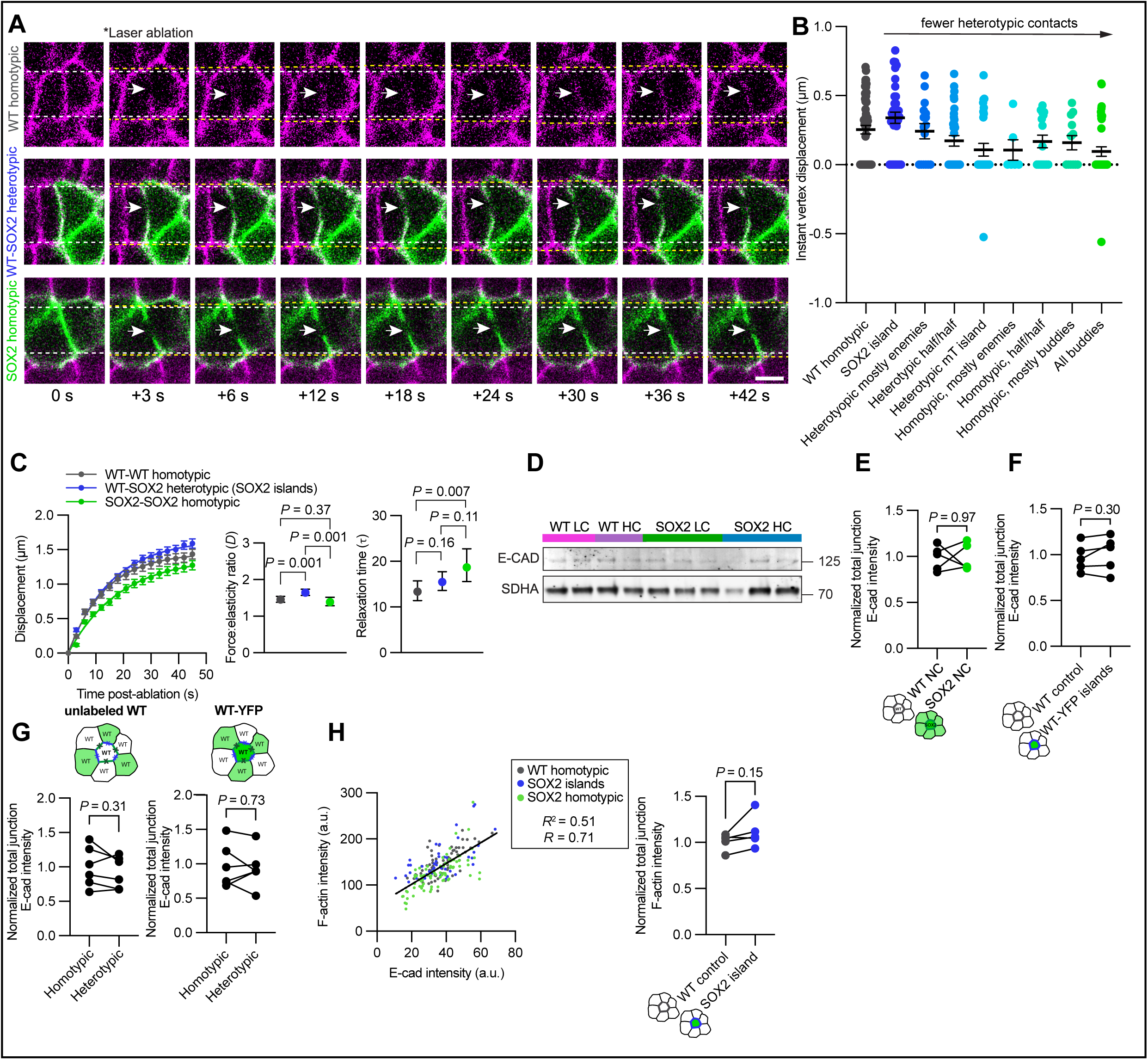
Laser ablation parameter fitting, F-actin correlation, and controls supporting heterotypic SOX2-WT interface characterisation. **A)** Representative laser-induced ablations at WT homotypic **(top row)**, WT-SOX2 heterotypic **(middle row)**, and SOX2 homotypic **(bottom row)** borders in E14.5 embryos. White dashed lines indicate the initial vertex locations in the time frame immediately preceding ablation (0 s); yellow dashed lines indicate the displaced vertex locations at each frame shown; white arrows denote the location of the cut/ablated border. **B)** Quantification of instant vertex displacement following laser ablation in E14.5 embryos across junctions in neighborhood conformations in decreasing heterotypic degree. Two vertices are plotted per junction. *n* = 17 WT homotypic and 14 WT island heterotypic junctions (across ≥ 8 embryos, ≥ 3 litters total); two-tailed unpaired *t*-test with Welch’s correction. **C)** Laser ablation of WT homotypic, WT-SOX2 heterotypic, and SOX2 homotypic borders within E14.5 mosaic skin. **Left,** displacement of individual border vertices plotted against time (3 sec intervals) where *t =* 0 denotes time immediately preceding ablation. Estimated force:elasticity ratios (*D*; **middle**) and relaxation time (tau; **right**) extracted from fit of laser ablation data for each junctional class, plotted with 95% confidence intervals. *n* = 36 WT-WT homotypic; 30 WT-SOX2 heterotypic islands; 42 SOX2-SOX2 homotypic junctions (across ≥ 15 embryos, ≥ 8 litters total). **D)** Western blot of E-cadherin between sorted E14.5 LC WT, HC WT, LC SOX2, and HC SOX2 EpSCs. Quantification displayed in **Fig. 2D**. **E)** Quantification of E-cad intensity across all junctions between exclusively non-competitive WT and SOX2 EpSCs. Values are normalized to average control WT across all embryos and litters. *n* ≥ 2 embryos per litter; 2 litters total; two-tailed paired *t*-test. **F)** Quantification of E-cad intensity across all junctions between non-competitive WT and WT-YFP EpSCs. Values are normalized to average control WT across all embryos and litters. *n* ≥ 2 embryos per litter; 2 litters total; two-tailed paired *t*-test. **G)** Heterotypic and homotypic border analysis of E-cad intensity in WT-YFP control mosaics between unlabelled WT and labelled WT EpSCs. *n* ≥ 2 embryos per litter; 2 litters total; two-tailed paired *t*-test. **H) Left,** correlation of raw E-cad and F-actin intensities of total junctions in non-competitive WT (homotypic) controls, SOX2 islands (competitive, heterotypic), and non-competitive SOX2 (homotypic) EpSCs. *n ≥* 10 cells per condition per embryo, ≥ 2 embryos per litter, 2 litters total. Non-linear regression performed on pooled measurements. **Right,** quantification of F-actin intensity across all junctions between non-competitive WT and SOX2 islands. Values are normalized to average control WT across all embryos and litters. *n* ≥ 2 embryos per litter; 2 litters total; two-tailed paired *t*-test. All data, except for the 95% CIs, are shown as mean ± s.e.m. Scale bar = 10 µm.

**Supplemental Figure 4.**
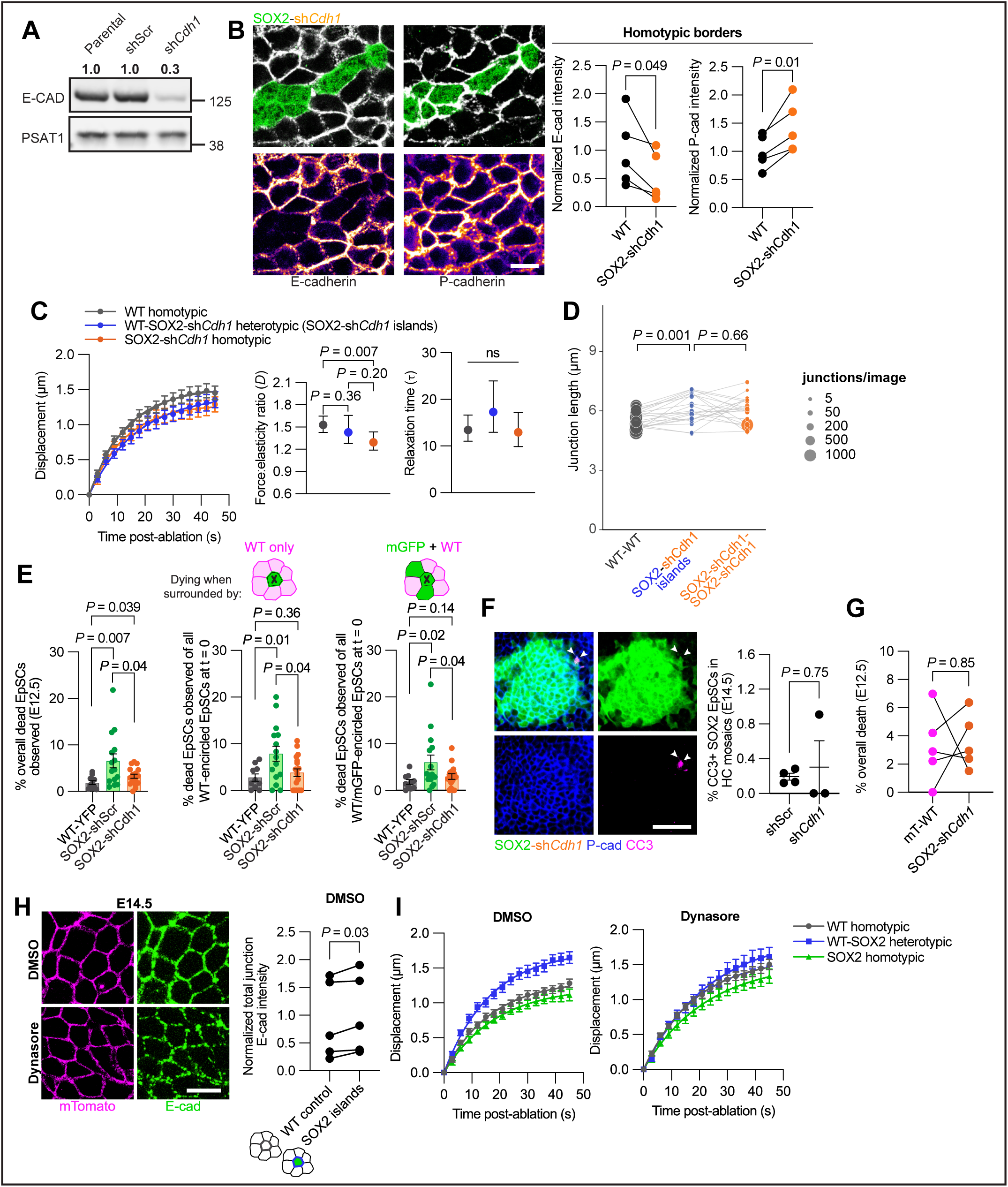
Consequences of Cdh1 knockdown and endocytic inhibition in SOX2 EpSCs. **A)** Western blot analysis of WT keratinocytes transduced with shScr and sh*Cdh1* hairpins. Quantifications shown above blot. **B) Left,** representative whole-mount maximum projection immunofluorescence image of E14.5 epidermis mosaically populated by SOX2^sh*Cdh1*^ EpSCs and stained for E-cad **(left)** and P-cad **(right)**. **Right,** quantification of normalized E-cad **(left)** and P-cad **(right)** at homotypic WT and SOX2^sh*Cdh1*^ borders. **C)** Laser ablation of WT homotypic, WT-SOX2^sh*Cdh1*^ heterotypic, and SOX2^sh*Cdh1*^ homotypic borders within E14.5 mosaic embryos. **Left,** displacement of individual border vertices plotted against time (3 sec intervals) where *t =* 0 denotes time immediately preceding ablation. Estimated force:elasticity ratios (*D*; **middle**) and relaxation time (tau; **right**) extracted from fit of laser ablation data for each junctional class, plotted with 95% confidence intervals. *n* = 15 WT-WT homotypic; 16 WT-SOX2^sh*Cdh1*^ heterotypic islands; 16 SOX2^sh*Cdh1*^-SOX2^sh*Cdh1*^ homotypic junctions (across ≥ 7 embryos, ≥ 4 litters total). **D)** Quantification of individual lengths between WT-WT homotypic, WT-SOX2^sh*Cdh1*^ heterotypic island, and SOX2^sh*Cdh1*^-SOX2^sh*Cdh1*^ homotypic junctions. Each datapoint represents measurements extracted from the average value of all cells in each category within a single field of view/image; lines connect matched values from within the same image. *n* = 19 images across 5 embryos from 2 litters. Statistical comparisons were made via a paired t-test with x3 Bonferroni correction. **E)** Quantifications of death events from E12.5 live imaging experiments between WT-YFP, SOX2^shScr^ (re-plotted from **Fig. 1, F and H**), and SOX2^sh*Cdh1*^ EpSCs. Shown are percentages of EpSC deaths for: **left**, total; **middle**, those encircled by WT EpSCs only; and **right**, those surrounded by a mixed combination of WT and SOX2 neighbors. *n =* 1-5 FOVs per embryo; 3-5 embryos per condition; two-tailed unpaired *t*-test with Welch’s correction for each statistical comparison. **F) Left,** representative whole-mount maximum projection immunofluorescence image of E14.5 epidermis populated with SOX2^sh*Cdh1*^ EpSCs at high-coverage. White arrowheads denote dying clones, which are primarily in contact with WT neighbors but not within the bulk of the clone. **Right,** quantification of CC3+ SOX2 EpSCs at high-coverage comparing shScr and sh*Cdh1. n* ≥ 3 embryos across 2 litters; two-tailed unpaired *t*-test with Welch’s correction. **G)** Quantification of death events in E12.5 live-imaged embryonic epidermis between mT-WT and SOX2^sh*Cdh1*^ clones. *n* = 5 high-coverage FOVs from 3 embryos; two-tailed paired *t*-test. **H) Left,** representative whole-mount maximum projection immunofluorescence image of E14.5 epidermis treated with either DMSO **(top)** or Dynasore **(bottom). Right,** quantification of E-cad intensity across all junctions between WT control and SOX2 island EpSCs in DMSO-treated E14.5 embryos. Quantification in Dynasore-treated embryos displayed in **Fig. 2K**. All values are normalized to average control WT in DMSO treatment across all embryos and litters. *n* ≥ 2 embryos per litter; 2 litters total; two-tailed paired *t*-test. **I)** Laser ablation of WT homotypic, WT-SOX2 heterotypic, and SOX2 homotypic borders within DMSO or Dynasore-treated E14.5 embryos. **Left,** displacement of individual border vertices plotted against time (3 sec intervals) where *t =* 0 denotes time immediately preceding ablation. Estimated force:elasticity ratios (*D*; **middle**) and relaxation time (tau; **right**) extracted from fit of laser ablation data for each junctional class, plotted with 95% confidence intervals. For DMSO condition, *n* = 14 WT-WT homotypic; 12 WT-SOX2 heterotypic; 13 SOX2-SOX2 homotypic junctions (across ≥ 5 embryos, 3 litters total). For Dynasore, *n* = 17 WT-WT homotypic; 16 WT-SOX2 heterotypic; 15 SOX2-SOX2 homotypic junctions (across ≥ 5 embryos, 3 litters total) All data shown as mean ± s.e.m. Scale bars = 10 µm **(B, H)**; 50 µm **(F)**.

**Supplemental Figure 5.**
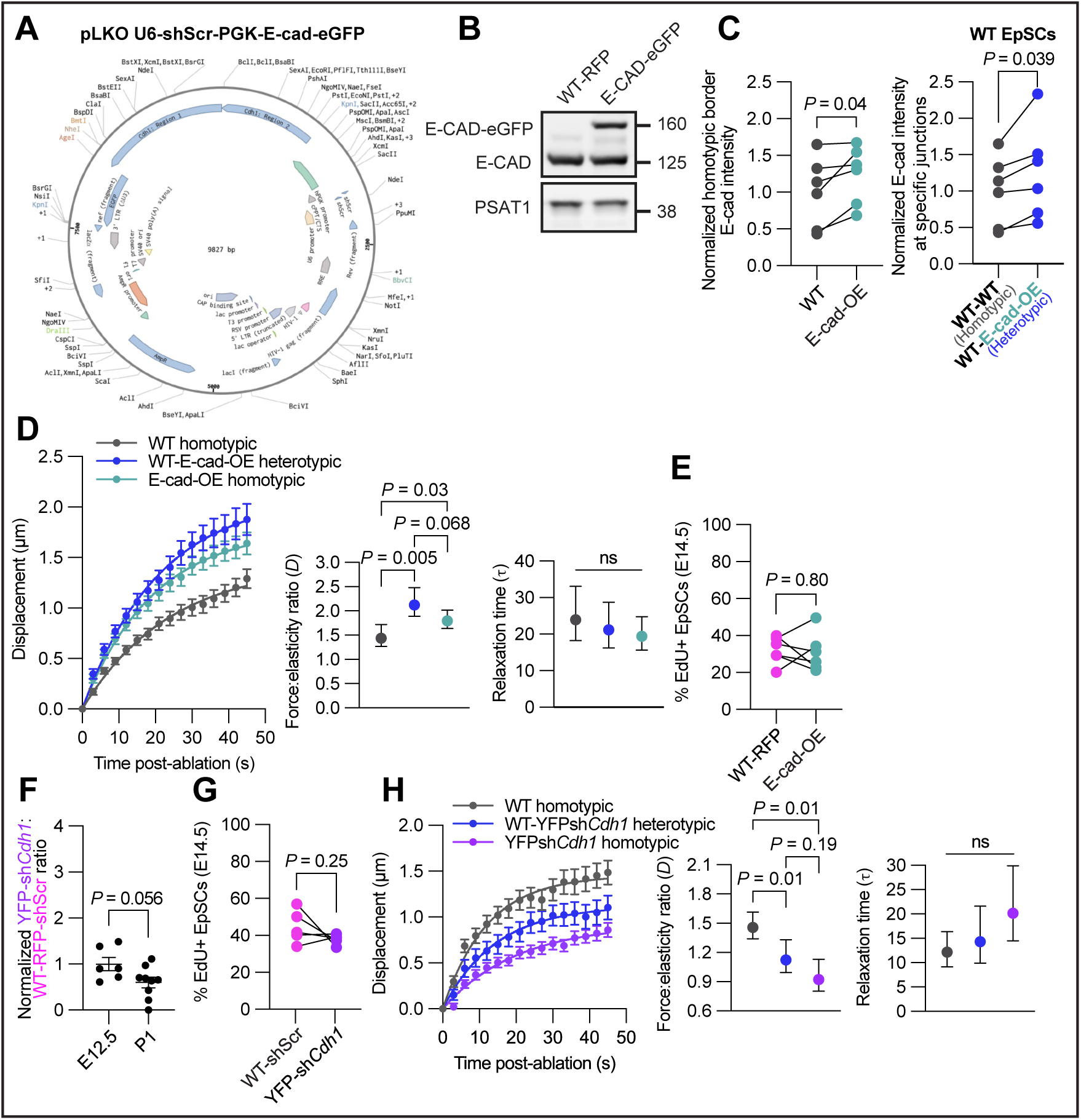
Validation and supporting analyses for mosaic E-cadherin gain- and loss-of-function experiments. **A)** Vector map for E-cadherin over-expression construct. **B)** Western blot analysis of keratinocytes transduced with WT-RFP and E-cad^OE^ constructs. **C)** Left, quantification of normalized E-cad intensity at the homotypic borders between WT controls and E-cad^OE^ EpSCs. **Right**, quantification of normalized E-cad intensity between heterotypic and homotypic borders within WT EpSCs neighboring E-cad^OE^ EpSCs. Normalisation is performed using the average WT homotypic control across all embryos and litters. **D)** Laser ablation of WT homotypic, WT-E-cad^OE^ heterotypic, and E-cad^OE^ homotypic borders within E14.5 embryos. **Left,** displacement of individual border vertices plotted against time (3 sec intervals) where *t =* 0 denotes time immediately preceding ablation. Estimated force:elasticity ratios (*D*; **middle**) and relaxation time (tau; **right**) extracted from fit of laser ablation data for each junctional class, plotted with 95% confidence intervals. *n* = 13 WT-WT homotypic; 17 WT-E-cad^OE^ heterotypic islands; 12 E-cad^OE^-E-cad^OE^ homotypic junctions (across ≥ 4 embryos, ≥ 3 litters total). **E)** EdU incorporation assay reveals that proliferation is comparable between E-cad^OE^ and WT EpSCs at E14.5. *n* ≥ 2 embryos per litter; 2 litters per time point; two-tailed paired *t*-test. **F)** Comparative growth assay between WT and YFP^sh*Cdh1*^ EpSCs during embryonic development. *n* ≥ 2 embryos per litter; 2 litters per time point; two-tailed unpaired *t*-test with Welch’s correction. **G)** EdU incorporation assay reveals that proliferation is comparable between YFP^sh*Cdh1*^ and WT EpSCs at E14.5. *n* ≥ 2 embryos per litter; 2 litters per time point; two-tailed paired *t*-test. **H)** Laser ablation of WT homotypic, WT-YFP^sh*Cdh1*^ heterotypic, and YFP^sh*Cdh1*^ homotypic borders within E14.5 embryos. **Left,** displacement of individual border vertices plotted against time (3 sec intervals) where *t =* 0 denotes time immediately preceding ablation. Estimated force:elasticity ratios (*D*; **middle**) and relaxation time (tau; **right**) extracted from fit of laser ablation data for each junctional class, plotted with 95% confidence intervals. *n* = 14 of all 3 junctions (across ≥ 8 embryos, 2 litters total). All data shown as mean ± s.e.m.

**Supplemental Figure 6.**
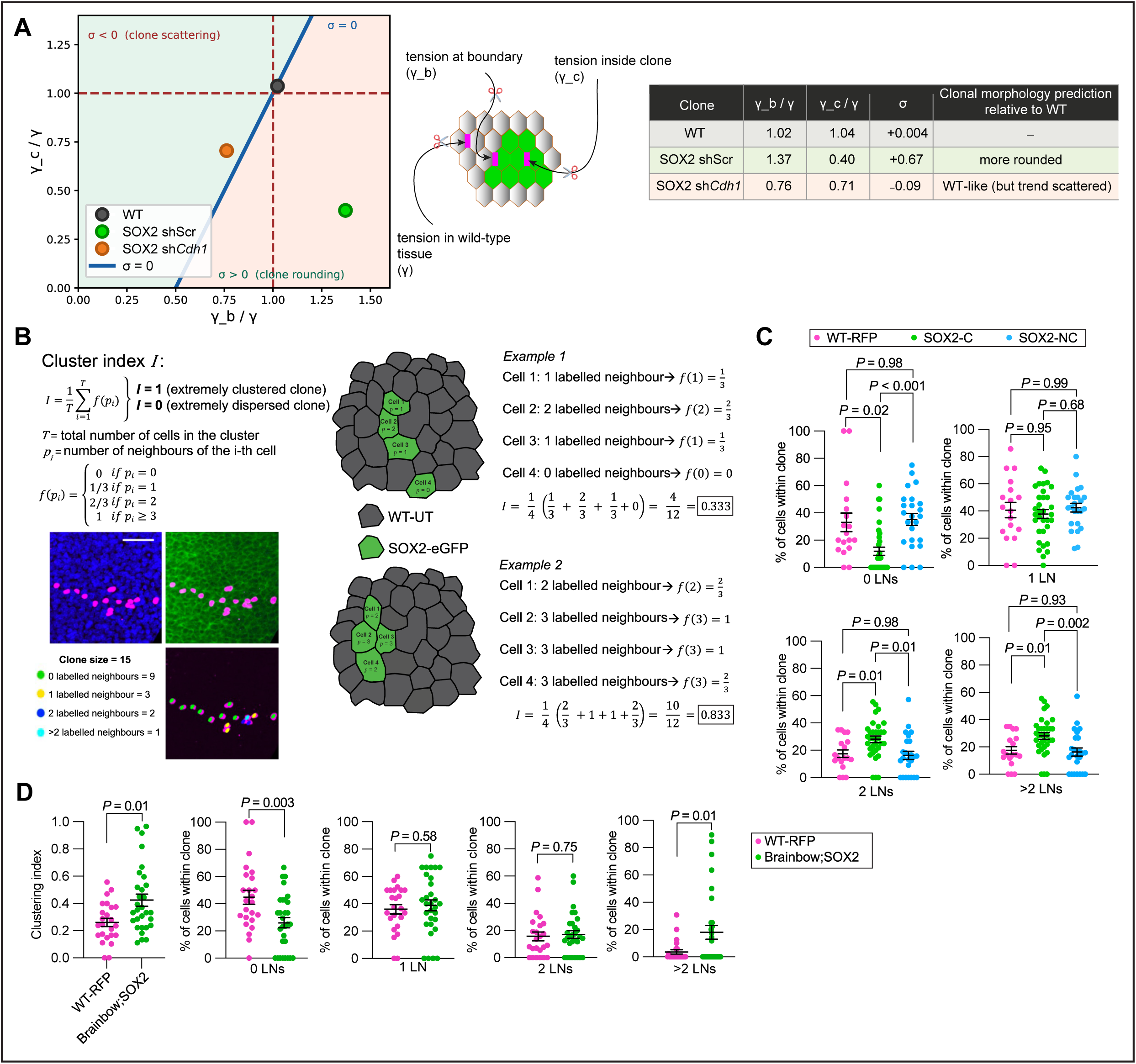
Quantitative analysis of clustering in validated clones. **A)** Phase diagram predicting clonal morphology from tension distributions. Position in (γ_b/γ, γ_c/γ) space — boundary tension and intra-clonal tension, each normalized to wild-type tissue tension — predicts whether clones scatter or round up. σ represents the net interfacial energy parameter governing clonal morphology: σ < 0 favours clone scattering (green region), σ > 0 favours clone rounding (peach region), and the line σ = 0 (blue) marks the boundary between these regimes. Schematic at right defines γ_b (tension at clonal boundary), γ_c (tension inside clone), and γ (tension in surrounding wild-type tissue). Data points indicate measured (γ_b/γ, γ_c/γ) values for WT (grey), SOX2 shScr (green), and SOX2 sh*Cdh1* (orange) clones. Table summarizes values and predicted clonal morphology relative to WT. Phase diagram adapted from Bosveld et al (reference 47). **B)** Definition of the cluster index *I*, where *T* is the total number of cells in the clone, *p_i_* is the number of directly adjacent neighbors of cell *i*, and *f*(*p_i_*) is a weighting function reflecting the contribution of each cell based on its local neighborhood. Two example clones illustrate how the cluster index is calculated. Each cell’s number of direct neighbors is shown, and the corresponding *f*(*p_i_*) values are indicated. The resulting cluster index *I* for each example is shown next to the schematic. The first example represents a low-cohesiveness clone with sparse connectivity and a correspondingly low *I* value, while the second example shows a highly cohesive clone with dense local clustering and a correspondingly high *I* value. **C)** Expanded clonal dispersion assay comparing WT-RFP, competitive SOX2, and non-competitive SOX2 EpSC clones at E14.5. *n* = 17 WT clones (5 embryos across 2 litters); 33 competitive SOX2 clones (6 embryos across 2 litters); 23 non-competitive SOX2 clones (3 embryos across 2 litters); Brown-Forsythe and Welch ANOVA with Dunnett’s T3 multiple comparisons test. **D)** Clonal dispersion assay comparing Brainbow;SOX2 and internal WT EpSC clones. LNs = labelled neighbors. *n* = 24 YFP clones (7 embryos across 2 litters); 30 SOX2 clones (7 embryos across 2 litters); two-tailed unpaired *t*-test with Welch’s correction. Scale bar = 50 µm.

**Supplemental Figure 7.**
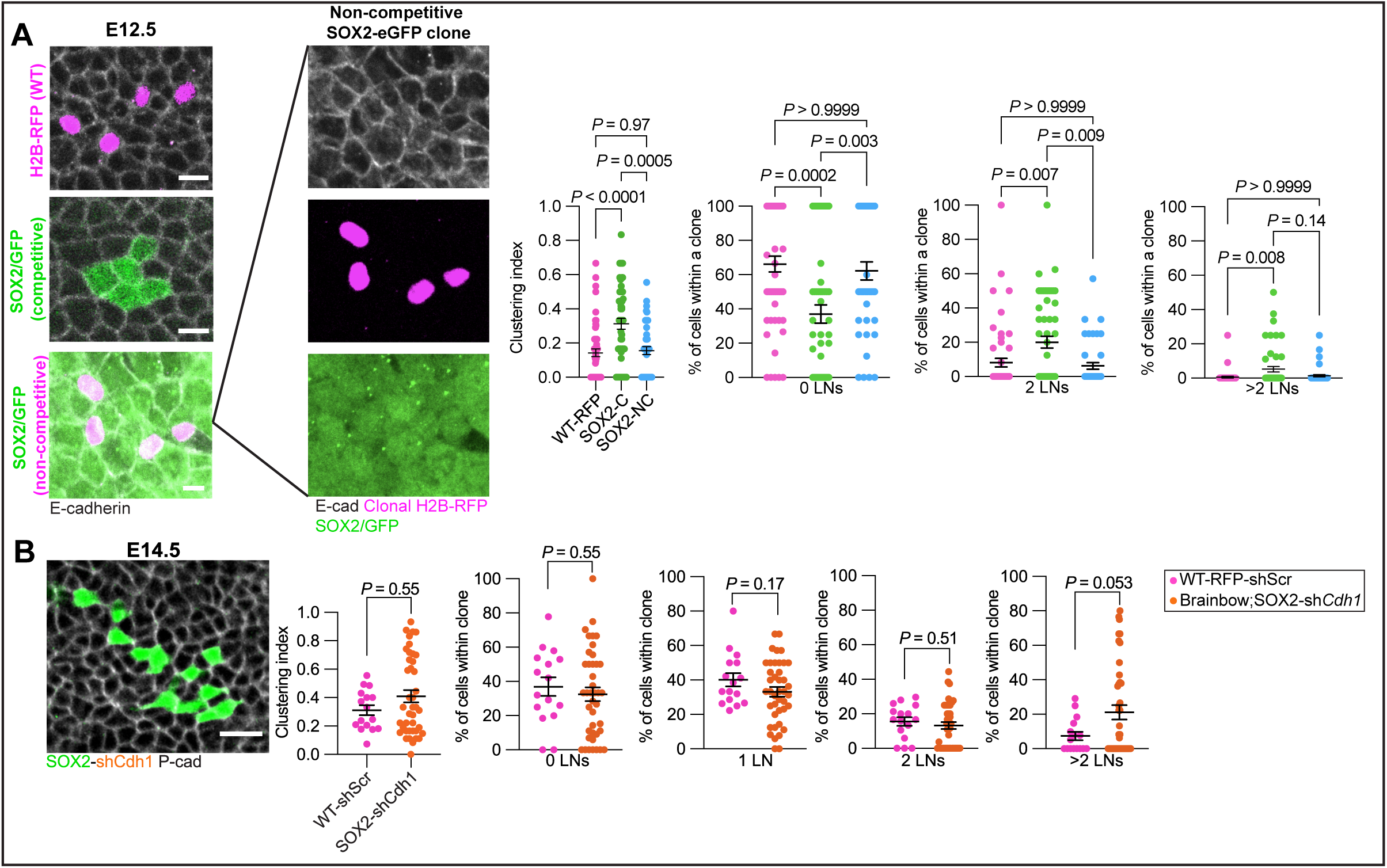
Quantitative analysis of clonal clustering in mosaic epidermis. **A)** Clonal dispersion assay comparing WT-RFP, competitive SOX2, and non-competitive SOX2 EpSC clones at E12.5. *n* = 54 WT clones (5 embryos across 2 litters); 52 competitive SOX2 clones (5 embryos across 2 litters); 43 non-competitive SOX2 clones (5 embryos across 2 litters); Brown-Forsythe and Welch ANOVA with Dunnett’s T3 multiple comparisons test. **B) Left,** representative immunofluorescence image revealing the dispersed nature of a SOX2^sh*Cdh1*^ EpSC clone in a competitive environment. Accompanying quantifications using clustering index demonstrate that clustering now more closely resembles WT EpSC behavior. *n =* 16 WT clones (from 3-6 embryos per litter; 2 litters); 41 SOX2^sh*Cdh1*^ clones (from 4 embryos per litter; 2 litters); two-tailed unpaired Mann-Whitney U test. **Right,** expanded clonal dispersion assay comparing SOX2^sh*Cdh1*^ and internal WT controls. *n =* 16 WT clones (from 3-6 embryos per litter; 2 litters); 41 SOX2^sh*Cdh1*^ clones (from 4 embryos per litter; 2 litters); two-tailed unpaired *t-*test. All data shown as mean ± s.e.m. Scale bar = 10 µm **(A)**, 20 µm **(B)**.

**Supplemental Figure 8.**
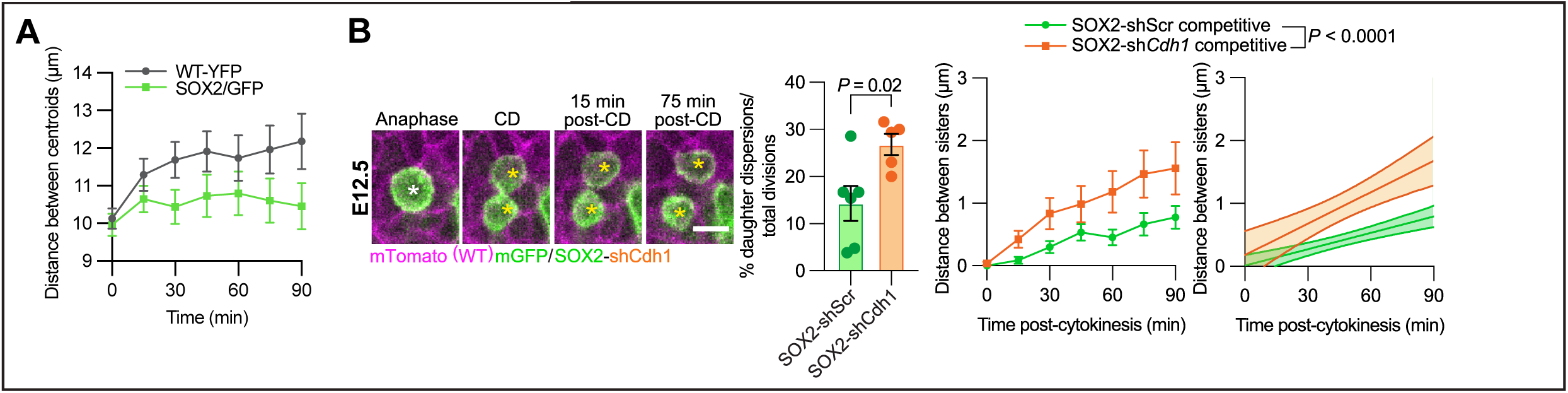
E-cadherin-dependent cohesion of SOX2 daughter EpSCs following mitosis. **A)** Quantification of clonal dispersion of E12.5 live-imaged EpSCs using centroid as focal point for analysis. Distance between daughter centroids was then measured over time. **B) Left,** representative E12.5 SOX2^sh*Cdh1*^ daughters disperse following cell division (CD). **Middle left,** quantification of percentages of SOX2^sh*Cdh1*^ clones whose daughters dispersed following division (SOX2^shScr^ data re-graphed from **Fig. 4B**). *n ≥* 5 embryos for each condition. **Middle right**, quantification of the distance between resulting daughter cells following division. **Right,** nonlinear regressions shown as 95% confidence intervals comparing the dispersion of SOX2^shScr^ and SOX2^sh*Cdh1*^ post-division. Statistics performed on nonlinear regressions. Note that SOX2^sh*Cdh1*^ EpSC clones dispersed more than SOX2^shScr^ EpSCs in competitive environments. *n ≥* 70 clones from 5 embryos per condition. All data shown as mean ± s.e.m. Scale bar = 10 µm.

**Supplemental Figure 9.**
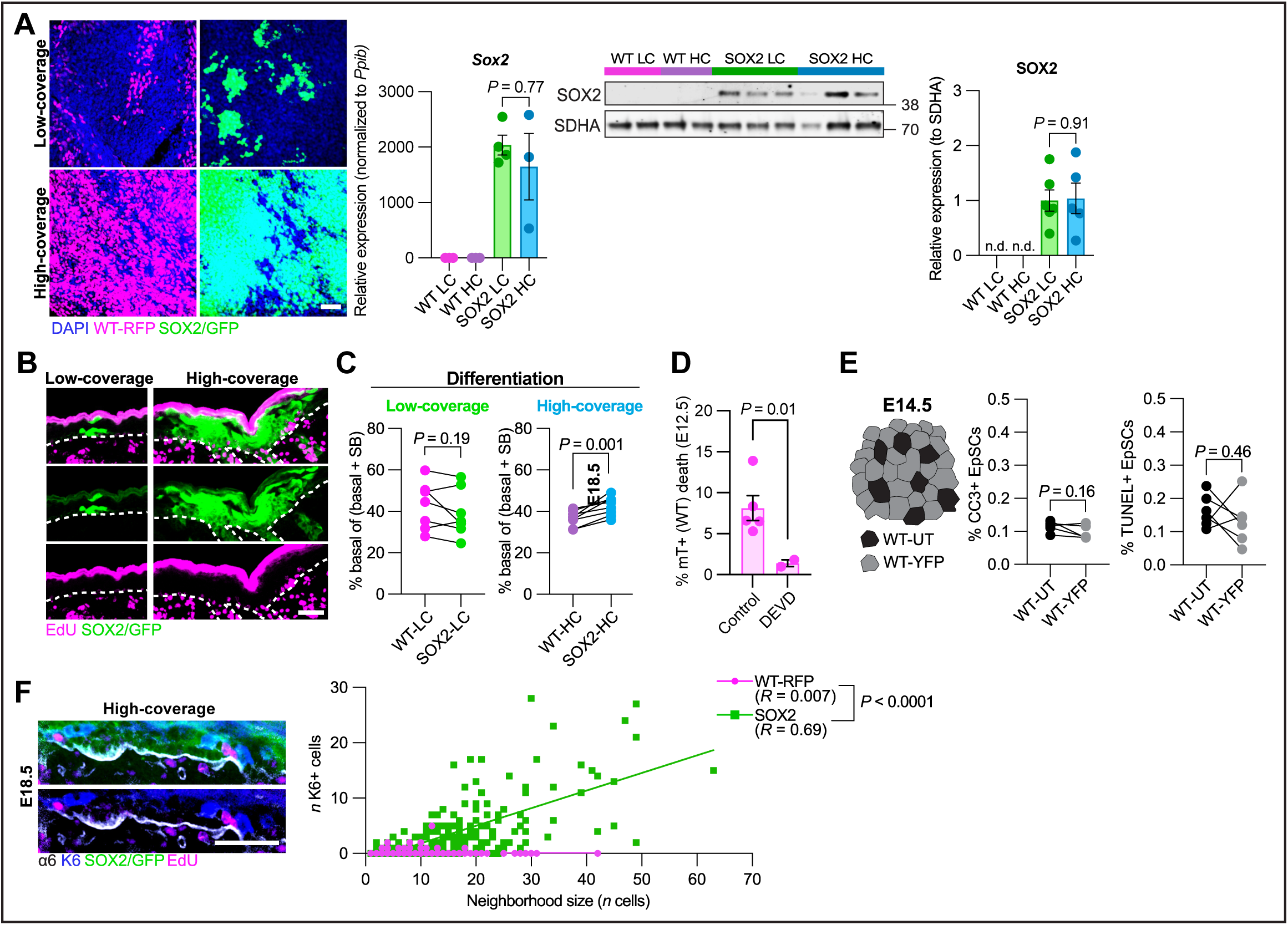
Cellular and molecular evidence for competitive regime switching in SOX2-dominated epidermis. **A) Left,** representative E14.5 whole-mount maximum projection immunofluorescence images of samples analyzed for SOX2 mRNA transcript levels via qPCR (**middle left)** and SOX2 protein via Western blot analysis (**middle right, right)** between low-coverage **(top)** and high-coverage **(bottom)** WT and SOX2 EpSCs. For qPCR, ordinary one-way ANOVA with Tukey’s multiple comparisons test. For Western blot, two-tailed unpaired *t*-test with Welch’s correction. **B)** Representative sagittal immunofluorescence images of SOX2 clones in LC and HC mosaics stained for EdU. White dashed line denotes epidermis-dermis boundary. Note that significant off-target signal accumulates in stratum corneum. **C)** Quantifications of basal layer retention in SOX2 EpSCs Basal-layer retention was measured as the percentage of basal cells within total epidermal cell populations (basal, B + suprabasal, SB). *n ≥* 2 embryos per condition per litter; 2 litters total; two-tailed paired *t-*test. **D)** Quantifications of dying mT WT EpSCs in HC SOX2 embryos treated with DMSO and DEVD. *n ≥* 2 FOVs per embryo, ≥ 2 embryos total; two-tailed unpaired *t-*test with Welch’s correction. **E)** Schematic of high-coverage WT-YFP EpSCs surrounding WT-untransduced (WT-UT) EpSCs. Quantifications reveal comparable CC3 and TUNEL labeling when WT EpSCs are surrounded by WT neighbors. *n ≥* 2 embryos per litter; 2 litters total; two-tailed paired *t-*test. **F) Left,** representative sagittal section of E18.5 epidermis where EdU+ SOX2 EpSCs in HC mosaics localize adjacent to K6+ SOX2 cells. **Right,** correlation analysis between neighborhood size and K6 expression within SOX2 and WT-RFP clones. *n ≥* 147 WT-RFP neighborhoods; 236 SOX2-eGFP neighborhoods (from ≥ 2 embryos per condition; 2 conditions per genotype per litter; 2 litters). All data shown as mean ± s.e.m. Scale bars = 50 µm.

**Supplemental Figure 10.**
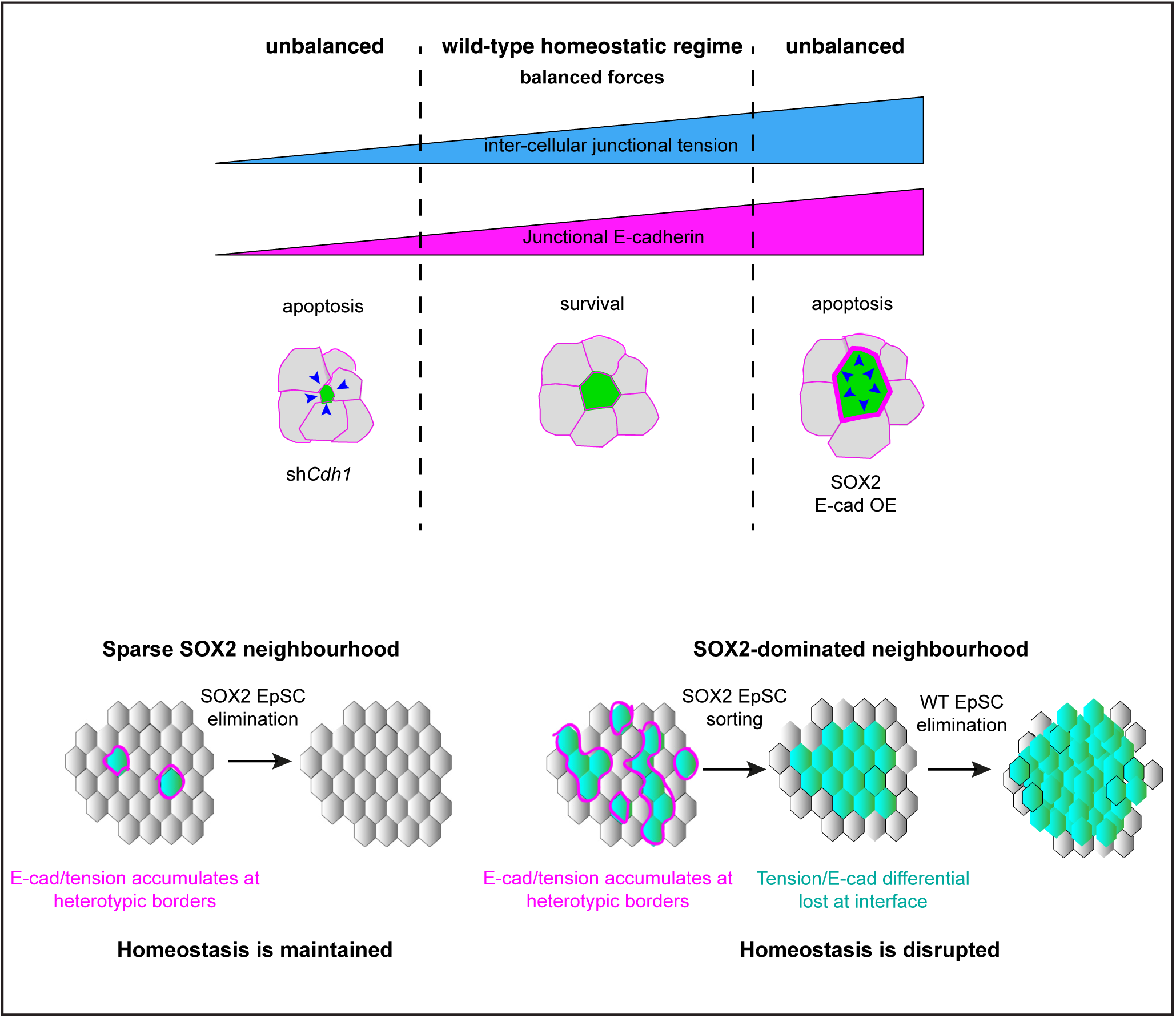
Bidirectional tension-based fitness control and the route to homeostatic collapse in mosaic epidermis.

## SUPPLEMENTARY INFORMATION

**Supplementary Video S1. Representative example of a SOX2 EpSC surrounded entirely by WT neighbors undergoing cell death.** Time-lapse imaging of the basal epidermis of an intact E12.5 embryo shows SOX2 EpSCs (green, mGFP+) are more likely to die dependent on contact with WT neighbors (magenta, mTomato+).

**Supplementary Video S2. Representative example of a SOX2 EpSC in contact with both SOX2 and WT neighbors undergoing cell death.** Time-lapse imaging of the basal epidermis of an intact E12.5 embryo shows SOX2 EpSCs (green, mGFP+) are more likely to die dependent on contact with WT neighbors (magenta, mTomato+).

**Supplementary Video S3. Representative example of a SOX2 EpSC surrounded entirely by SOX2 neighbors undergoing cell death.** Time-lapse imaging of the basal epidermis of an intact E12.5 embryo shows a rarely occurring example of a dying SOX2 EpSCs (green, mGFP+) that is only making cell-cell contacts with other SOX2 EpSCs.

**Supplementary Video S4. Representative example of laser ablation of a WT homotypic border.** Homotypic junctions shared between two neighboring mTomato+ WT EpSCs in the basal layer of E14.5 embryos were subject to ablation. Time-lapse imaging was simultaneously performed to record the initial position of the selected junctions (2 frames are shown pre-cut) and to allow for visualization of the tissue response, post-ablation. The laser pulse occurs at t=0 at which point junctional severing can be observed. The junctional vertices at either end of the cut junction retract and displace relative to their initial position. The rate and the extent of displacement is measured at each time interval (images were acquired every three seconds for 42 seconds post-ablation). WT homotypic borders displayed retraction behaviors that were distinct from WT-SOX heterotypic borders and SOX2 homotypic borders.

**Supplementary Video S5. Representative example of laser ablation of a WT-SOX2 heterotypic border.** Heterotypic junctions shared between neighboring mTomato+ WT and mGFP+ SOX2 EpSCs in the basal layer of E14.5 embryos were subject to ablation. Time-lapse imaging was simultaneously performed to record the initial position of the selected junctions (2 frames are shown pre-cut) and to allow for visualization of the tissue response, post-ablation. The laser pulse occurs at t=0 at which point junctional severing can be observed. The junctional vertices at either end of the cut junction retract and displace relative to their initial position. The rate and the extent of displacement is measured at each time interval (images were acquired every three seconds for 42 seconds post-ablation). WT-SOX heterotypic borders displayed retraction behaviors that were distinct from WT homotypic borders and SOX2 homotypic borders.

**Supplementary Video S6. Representative example of laser ablation of a SOX2 homotypic border.** Homotypic junctions shared between two neighboring mGFP+ SOX2 EpSCs in the basal layer of E14.5 embryos were subject to ablation. Time-lapse imaging was simultaneously performed to record the initial position of the selected junctions (2 frames are shown pre-cut) and to allow for visualization of the tissue response, post-ablation. The laser pulse occurs at t=0 at which point junctional severing can be observed. The junctional vertices at either end of the cut junction retract and displace relative to their initial position. The rate and the extent of displacement is measured at each time interval (images were acquired every three seconds for 42 seconds post-ablation). SOX2 homotypic borders displayed retraction behaviors that were distinct from WT-SOX heterotypic borders and WT homotypic borders.

**Supplementary Video S7. Representative example of WT daughter cells dispersing following cell division in a non-competitive environment.** Time-lapse imaging of the basal epidermis of an intact E12.5 *R26-lox-stop-lox-YFP* embryo shows a sparsely labelled WT-YFP EpSC (green, mGFP+) undergoing mitosis beginning from anaphase until 75 minutes post division. Magenta, mTomato+ EpSCs are also WT. Daughter cells disperse from one another and become interspersed with clonally distinct WT neighbors (magenta, mTomato+).

**Supplementary Video S8. Example of SOX2 daughter cells dispersing following cell division in a competitive environment.** Time-lapse imaging of the basal epidermis of an intact E12.5 *R26-lox-stop-lox-Sox2-IRES-eGFP* embryo shows a sparsely labelled SOX2 EpSC (green, mGFP+) undergoing mitosis beginning from anaphase until 75 minutes post-division. Daughter cells remain in contact with one another and are not interspersed with WT neighbors (magenta, mTomato+).

**Supplementary Video S9. Representative example of SOX2 daughter cells dispersing following cell division in a non-competitive environment.** Time-lapse imaging of the basal epidermis of an intact E12.5 *R26-lox-stop-lox-Sox2-IRES-eGFP* embryo shows a SOX2 EpSC (green, mGFP+) in a non-competitive, SOX2-dominated environment undergoing mitosis beginning from anaphase until 75 minutes post division. Daughter cells disperse from one another and intermingle with other clonally distinct mGFP+ SOX2 EpSCs. The dividing cell and its daughters have no WT neighbors (magenta, mTomato+) present in its immediate vicinity.

**Supplementary Video S10. Representative example of SOX2*^shCdh1^* daughter cells dispersing following cell division in a competitive environment.** Time-lapse imaging of the basal epidermis of an intact E12.5 *R26-lox-stop-lox-Sox2-IRES-eGFP* embryo shows a sparsely labelled SOX2*^shCdh1^* EpSC (green, mGFP+) undergoing mitosis beginning from anaphase until 75 minutes post division. In contrast to SOX2*^shScr^* controls, daughter cells disperse from one another and become interspersed with clonally distinct WT neighbors (magenta, mTomato+).

**Supplementary Video S11. Representative example of a WT EpSC surrounded entirely by SOX2 neighbors undergoing cell death.** Time-lapse imaging of the basal epidermis of an intact E12.5 embryo shows when neighborhoods are dominated by SOX2 EpSCs (green, mGFP+), WT EpSCs (magenta, mTomato+) are more likely to undergo elimination.

